# BetaII-Spectrin Gaps and Patches Emerge from the Patterned Assembly of the Actin/Spectrin Membrane Skeleton in Human Motor Neuron Axons

**DOI:** 10.1101/2025.05.09.653215

**Authors:** Nahir Guadalupe Gazal, Maria Jose Castellanos-Montiel, Guillermina Bruno, Anna Kristina Franco-Flores, Sarah Lépine, Lale Gursu, Ghazal Haghi, Gilles Maussion, Wolfgang E. Reintsch, Fernando D. Stefani, Agustín Anastasía, Mariano Bisbal, Ezequiel Axel Gorostiza, Thomas M. Durcan, Nicolás Unsain

## Abstract

The actin/spectrin membrane-associated periodic skeleton (MPS) is a cytoskeletal structure that supports axonal integrity and function. Lower spinal motor neurons (MNs) are characterized by exceptionally long axons and are particularly susceptible to degeneration in a wide range of hereditary neuromuscular disorders, including amyotrophic lateral sclerosis. Using confocal and super-resolution imaging, we characterized the spatial distribution βII-spectrin and the assembly pattern of the MPS in human MN axons derived from induced pluripotent stem cells (iPSCs). We discovered a striking gap-and-patch pattern in the medial axon, where sharply demarcated βII-spectrin gaps alternate with patches containing a well-organized MPS. The pattern is acutely induced by the kinase inhibitor staurosporine and pharmacological inhibition of actin polymerization prevents patch formation, indicating a requirement for actin nucleation in MPS assembly. Our data supports a model in which spectrin incorporation into nascent MPS patches depletes neighboring regions, producing long-range gaps-and-patches patterns.

## Introduction

The actin/spectrin <u>m</u>embrane-associated <u>p</u>eriodic skeleton (MPS) consists of a periodic arrangement of F-actin rings separated by ~185 nm α/β-spectrin tetramer spacers, and is present in the axons and dendrites of all neuronal types examined across diverse animals species (Unsain, Stefani, et al., 2018; Leterrier, 2021). Since its discovery in 2013 (Xu et al., 2013), the MPS has been identified in neurons from the peripheral and central nervous systems of worms, flies, rodents, and humans (D’Este et al., 2016; He et al., 2016). Its phylogenetic conservation is consistent with the fundamental neuronal functions in which it has been implicated. For example, a spectrin-dependent skeleton protects axons from breaking during normal animal movements or under acute stress (Hammarlund et al., 2007; Krieg et al., 2017; Dubey et al., 2020). Additionally, the MPS organizes signaling platforms through the recruitment of signaling receptors (Zhou et al., 2019) and impacts membrane protein diffusion (Albrecht et al., 2016; Rentsch et al., 2024). It can also assemble with myosin to form functional actomyosin complexes, which in turn dynamically alter axon caliber and tension (Berger et al., 2018; Costa et al., 2020; T. Wang et al., 2020). Furthermore, MPS remodeling has been found to contribute towards axonal breakdown during developmental axon degeneration (Unsain, Bordenave, et al., 2018; G. Wang et al., 2019), for a review, see (Costa & Sousa, 2021). Taken together, the evidence suggests that the MPS is critical for preserving the function and structural stability of axons.

Lower spinal motor neurons (MNs) are characterized by exceptionally long axons, which can extend thousands of times the diameter of the cell soma. Their somas reside in the anterior horn of the spinal cord, receiving input from upper motor neurons and projecting axons through peripheral nerves to innervate skeletal muscles, enabling voluntary movement (Stifani, 2014). Motor neuron axons are particularly susceptible to structural instability and degeneration in a wide range of hereditary neuromuscular disorders, including amyotrophic lateral sclerosis (ALS). ALS is a neurodegenerative disease characterized by the progressive loss of MNs from the brain cortex, brainstem and spinal cord. About 90% of ALS cases are sporadic (sALS) while the remaining cases are familial (fALS) (Younger & Brown, 2023). Over 30 genes have been implicated in fALS. Among these, FUS (fused in sarcoma), TARDBP (transactive response DNA-binding protein) and SOD1 (Cu/Zn superoxide dismutase 1), account for approximately 20% of fALS, and have also been described in about 5% of sALS (Akçimen et al., 2023). In spite of their clinical relevance, a detailed description of βII-spectrin distribution and MPS organization in MN axons is still incomplete.

Spectrin functions as a symmetric hetero tetramer. Dimeric spectrin is formed by the lateral association of α and β monomers in a head-to-tail fashion. Dimers then associate in a head-to-head formation to produce the ~185 nm-long tetramer. Neurons express a single isoform of α-spectrin (αII) but multiple β-spectrin isoforms (βII, βIII, and βIV), which exhibit distinct subcellular localizations. βII-spectrin predominates in the axonal shaft, βIII-spectrin is enriched in the soma and dendrites, while βIV-spectrin is the exclusive β isoform at the axon initial segment and the node of Ranvier. The MPS is absent in nascent, immature neurites and assembles only after initial neurite extension. Most studies on MPS assembly have been conducted in cultured rat hippocampal neurons, where it was shown that during axon extension the MPS is better organized proximal to the soma, with a seemingly continuous proximal-to-distal gradient of organization, with no evident periodic distribution in the most distal portions. After around 10 days *in vitro* (DIV), on the other hand, all axonal portions reach a high degree of organization of the MPS (Zhong et al., 2014; Barabas et al., 2017). Recently, it was proposed that this continuous MPS organization arises from the coalescence of discontinuous “patches” of incomplete MPS units that originate in the distal axon and migrate proximally (Hofmann et al., 2022). These patches appear to form from spectrin tetramers accumulating in the growth cone, which then assemble into an incomplete periodic structure. The transport and assembly of spectrin into the MPS have also been studied in mature neurons *in situ* in *Caenorhabditis elegans* (Glomb et al., 2023). In this animal model, authors showed that spectrin is actively transported to distal axonal regions via kinesin motors, and that proper MPS assembly and maintenance requires a finely balanced spectrin supply. However, the precise mechanisms governing MPS formation in extending neurites and whether different neuronal types exhibit unique assembly patterns remain poorly understood.

In this work, we investigated βII-spectrin distribution and MPS organization and assembly dynamics in human induced pluripotent stem cell (iPSC)-derived MNs (Deneault et al., 2022). Noteworthy, we observed that βII-spectrin and αII-spectrin in distinct axonal sections exhibited sharp, periodic interruptions in their otherwise continuous distribution. These “gaps” were interspersed with “patches” bearing a well organized MPS, forming a distinct pattern along the axon. The occurrence of axonal sections with βII-spectrin gaps was unrelated to cell stress or caspase/calpain activity, but could be notably induced by the broad kinase inhibitor staurosporine. Using stimulated emission depletion (STED) nanoscopy, we confirmed that the MPS was well preserved within the patches. Analysis of individual axons along their entire length revealed that gaps and patches predominantly occurred in the medial axon. In contrast, spectrin proximal to these regions was continuously distributed with normal MPS organization, whereas the distal region exhibited a sharp reduction in spectrin levels and a lack of an MPS organization. These findings imply that gap and patch formation are correlated: once βII-spectrin is incorporated into a nascent MPS structure, it stops diffusing and accumulates in patches, depleting free βII-spectrin in gaps. In addition, we demonstrated that actin nucleation plays a role in this process, as latrunculin-A treatment prevents the acute effect of staurosporine. Furthermore, we extended our analysis into cellular models of hereditary ALS to assess whether the MPS is similarly organized under pathological conditions. Our findings provide a detailed characterization of βII-spectrin distribution and MPS organization in human neurons and provide new insights into the dynamic assembly of this cytoskeletal structure.

## Results

### βII-spectrin is distributed in a proximal to distal gradient in human iPSC-derived MN axons

We initiated our study on the distribution and organization of βII-spectrin in axons of human MNs at the single cell level. At a cell density that sustains healthy development and maturation for 2 weeks, human iPSC-derived MN cultures develop dense axonal areas (Supp. Fig 1A). Although this “bulk culture” approach allows efficient sampling of axonal features, it precludes the tracking of individual axons. To unequivocally trace individual axons from the cell soma to the tip, we seeded cells at low density to obtain sparse neurons and axons. iPSC-derived MNs do not develop properly in low density cultures and tend to degenerate within the first few days in culture. To overcome this limitation, cells were seeded at high density in the corner of a tilted petri dish and allowed to attach for 2 hrs (Supp. Fig 1B). After cell attachment, culture dishes were placed flat for 2 weeks under standard conditions. As a result, cells became distributed across the coverslip in a density gradient (“Density gradient culture”, Supp. Fig 1B). Cells toward the middle of the coverslip are sufficiently spaced to allow individual axons to be unambiguously traced from the soma to the tip (Fig. 1C and Supp. Fig 1C). Using quantitative confocal microscopy (see Materials and Methods), we compared the intensities of βII-spectrin in the proximal, medial and distal portion of axons. To analyze all axons regardless of their length (ranging from 300 to 1000 μm) the first quarter of the axonal length was defined as the *proximal* segment, the following two quarters were grouped as the *medial* portion, and the final quarter was designated as *distal*. We found that βII-spectrin intensity was high in the proximal segment, dropping sharply in the medial region, and decreasing further in the distal region (Fig. 1A and B). Qualitatively, the distribution of βII-spectrin presented three noteworthy features: not enriched in axonal tips (which were invariably blunt), was enriched in occasional axonal enlargements and exhibits portions with a gap-and-patch pattern (inserts in Fig. 1C and Supp. Fig 1C). All axonal tips lacked an expanded growth cone, and there was no accumulation of βII-spectrin (Fig. 1D), in contrast to what is observed in rat hippocampal neurons cultured for 7 days (Suppl. Fig. 1D) (Hofmann et al., 2022). Half of the axons presented axonal enlargements enriched in βII-spectrin (Fig. 1E), with a tendency to localize in the medial and distal portions (Suppl. Fig. 1E). Interestingly, the gap-and-patch patterns were present in about a quarter of the axons examined and were further investigated in bulk cultures to enhance sampling efficiency.

**Figure 1.**
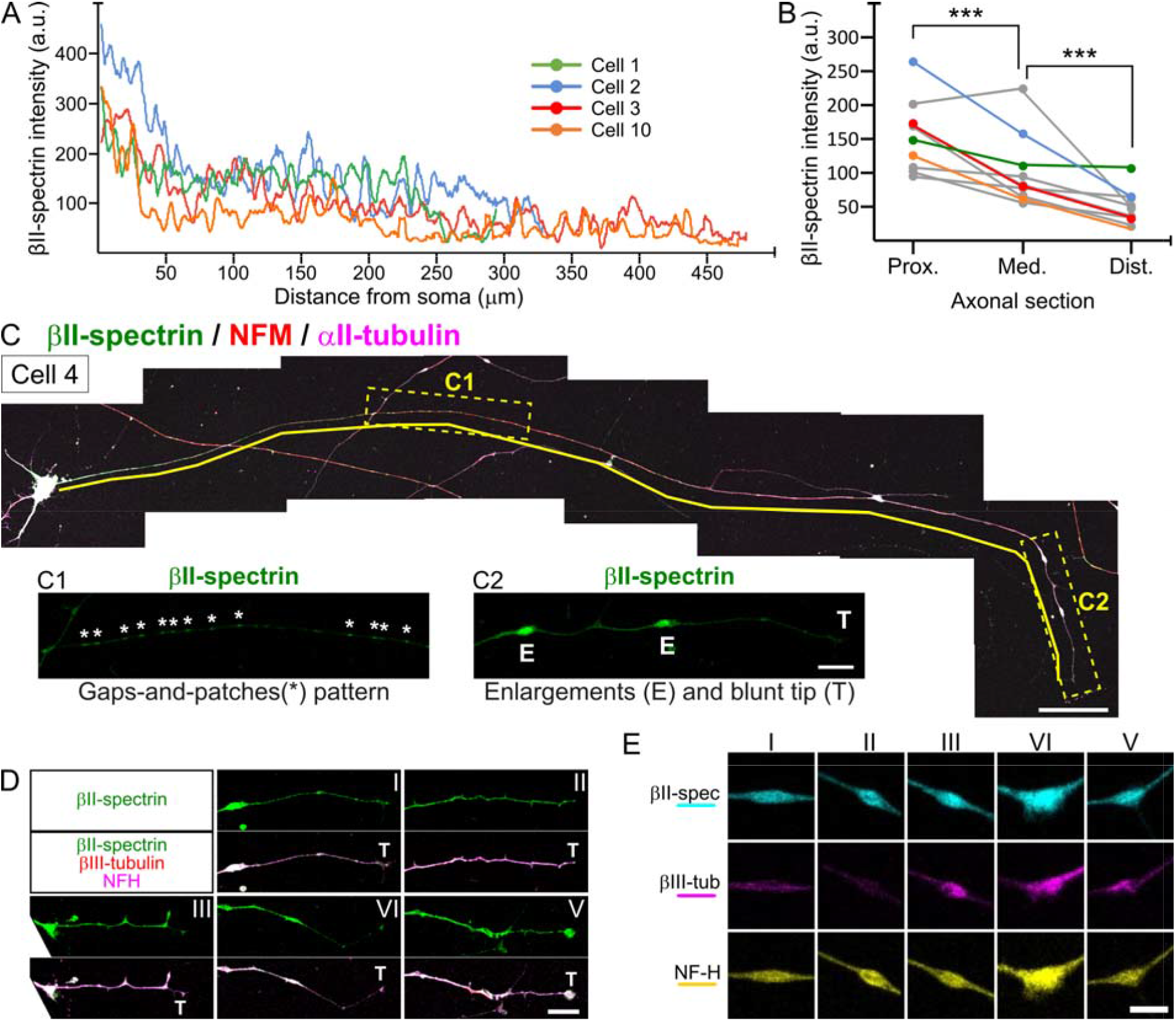
βII-spectrin distribution in human iPSC-derived MN axons. (**A**) Representative βII-spectrin intensity profiles along four individual axons. (**B**) Mean normalized βII-spectrin intensity in the proximal, medial and distal portion of 10 axons. Colored traces show the mean normalized values shown in panel A. ***: p<0.001. (**C**) Representative reconstruction of an individual axon, immunostained for βII-spectrin, NFM and αII-tubulin. Yellow line highlights the axon being followed. Scale bar = 50 μm. Inserts: C1 dashed rectangle highlights a gap-and-patch pattern (*: patches), whereas the C2 dashed rectangle highlights axonal enlargements (E) and the axonal tip (T). Scale bar = 10 μm. (**D**) Representative examples (I-V) of MN axonal tips (T) immunostained for βII-spectrin, βIII-tubulin and NFH. Scale bar = 10 μm. (**E**) Representative examples (I-V) of MN axonal enlargements immunostained for βII-spectrin, βIII-tubulin and NFH. Scale bar = 5 μm.

### Human iPSC-derived MNs exhibit long axonal sections with sharp interruptions in the distribution of βII-spectrin, creating a gap-and-patch pattern

Analyses of βII-spectrin distribution in bulk cultures revealed that a significant proportion of axonal segments exhibited pronounced irregularities, characterized by sharp interruptions, which we termed βII-spectrin “gaps” (βII-spec-gaps) (Fig. 2A). Gaps occurred along individual axons in consecutive sequences, forming a gap-and-patch pattern. When comparing the intensity of βII-spectrin in axons without gaps (“continuous”), within gaps, and within “patches” (βII-spectrin between gaps), we found that “gaps” had only 20% of the intensity observed in continuous sections, whereas patches presented a similar intensity to that of continuous sections (Suppl. Fig. 2A). Given these differences, we initially speculated that the gap-and-patch pattern found in certain segments was the end product of a local and spaced depletion of βII-spectrin.

**Figure 2.**
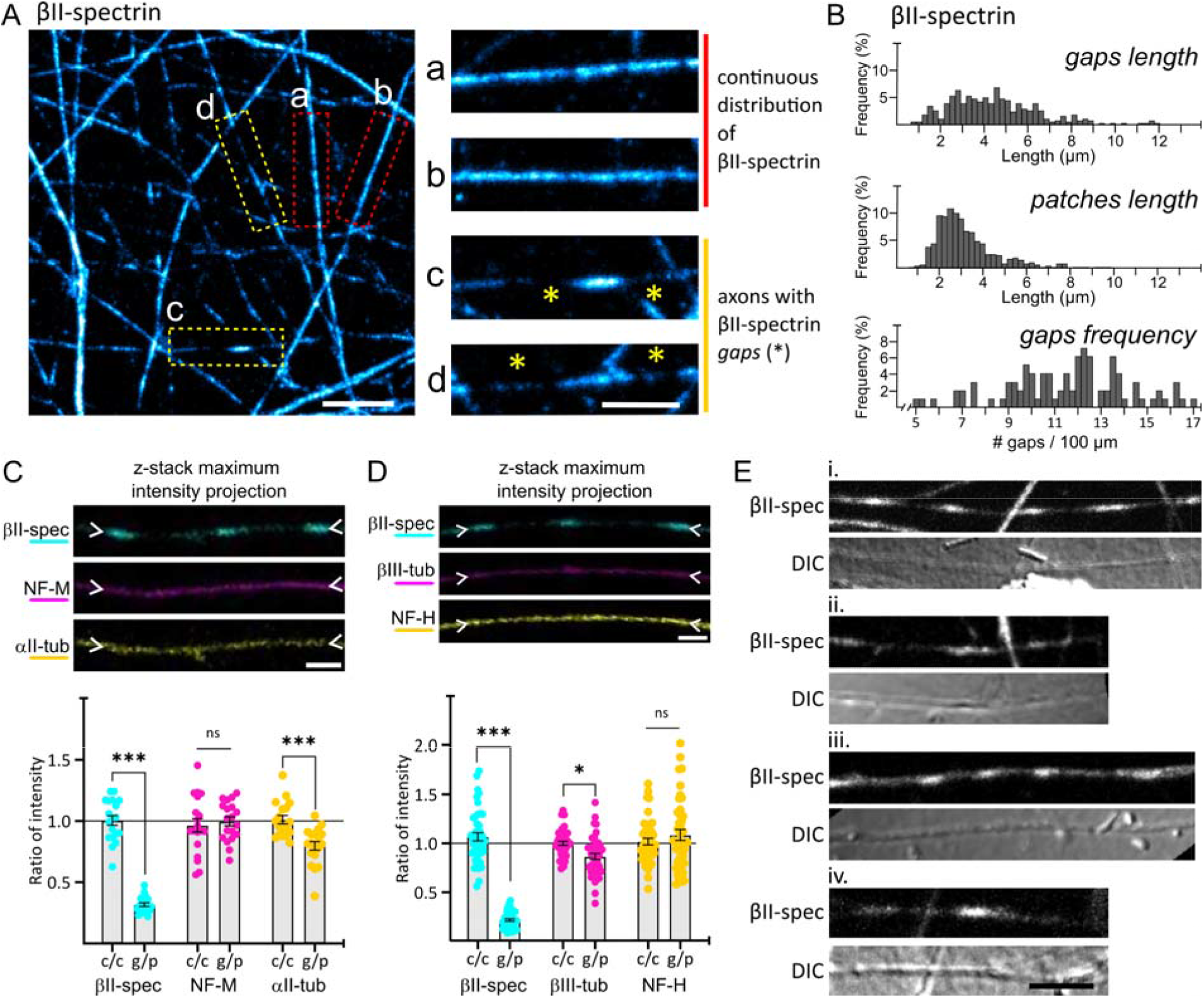
Motor neurons present sharp interruptions in the βII-spectrin lattice. (**A**) Confocal images of bulk cultures immunostained for βII-spectrin. Inserts a-c show examples of axons with a continuous distribution of βII-spectrin (a and b) and axons with sharp interruptions or “gaps” (asterisks in c and d). Scale bar = 10 μm (left panel) and 5 μm (zoom-in inserts). (**B**) Histograms showing percentual frequency of gaps length, patches length and gap frequency. (**C**) Top: Representative confocal images of a gap-and-patch pattern co-stained for βII-spectrin, Neurofilament M and □II-tubulin. Scale bar = 2.5 μm. Bottom: Quantification of intensity ratios. For axons with a continuous βII-spectrin distribution, the ratio of intensity between two regions 5 μm apart was measured (c/c). For axons with βII-spec-gaps, the ratio of intensity between a gap and its flanking patches was measured (g/p). Mean + SEM. T-test. ns: not significant; *: p<0.05; ***: p<0.001. (**D**) Top panel: Representative confocal images co-stained for βII-spectrin, βIII-tubulin and Neurofilament H. Scale bar = 2.5 μm. Lower panel: Quantification of intensity ratios. For axons with a continuous βII-spectrin distribution, the ratio of intensity between two regions 5 μm apart was measured (c/c). For axons with βII-spec-gaps, the ratio of intensity between a gap and its flanking patch was measured (g/p). Mean + SEM. T-test. ns: not significant; *: p<0.05; ***: p<0.001. (**E**) Confocal and *Differential Interference Contrast* images of axons immunostained for βII-spectrin with a gap-and-patch pattern. Scale bar = 5 μm.

These βII-spec-gaps had a mean length of 4.56 ± 2.50 μm (mean ± SD) and were spaced by βII-spectrin in “patches” (βII-spec-patches) with a mean length of 3.26 ± 1.46 μm (mean ± SD). In these axons, the gaps-and-patches pattern extended across the entire field of view in most cases, with a gap frequency of 12.13 ± 3.69 every 100 μm (mean ± SD, Fig. 2B). In our bulk cultures, in fields of view occupied only by axons, about 15% of axonal segments presented this pattern. A similar pattern of gaps and patches was observed when staining for αII-spectrin, strongly suggesting an absence of MPS in the gaps (Suppl. Fig. 2B and C).

To assess whether these gaps represented a gross loss of axonal material, we co-stained for constitutive components of the axonal cytoskeleton, with a focus on αII-tubulin, βIII-tubulin and neurofilament subunits heavy and medium (NF-H and NF-M) (Fig. 2C and D). The ratio of intensities between neighboring regions in axons without gaps (continuous axons) allowed us to estimate the variability of these protein distributions (c/c ratios; Fig 2C and D). Based on these findings, we next evaluated the intensity within gaps as a ratio to the neighboring patches. βII-spectrin intensity in gaps was approximately 20-30% of that found in patches (g/p ratios). The distribution of NF-M and NF-H did not change in gaps, while the distribution of αII- and βIII-tubulins showed a drop of 15% and 10%, respectively. (Fig. 2C and D). Using differential interference contrast (DIC) microscopy, no overt changes in axonal appearance or caliber were noticeable between βII-spec-gaps and the neighboring βII-spec-patches (Fig. 2E). Taking into account the continuity and persistence of neurofilament and tubulin isoforms within gaps, as well as the regular appearance of axons under DIC microscopy, we conclude that gaps do not represent axonal interruptions or a gross loss of axoplasm.

### Axonal segments bearing βII-spectrin gaps increase as a function of weeks in culture and by the acute treatment with staurosporine

To gain insight into the mechanisms underlying the formation of βII-spec-gaps, we next investigated conditions that might affect their occurrence. We first observed that the percentage of axonal segments with gaps increased as MNs mature in culture, from 1 to 3 weeks (Fig. 3A). Based on earlier observations, these pure MN cultures progressively coalesce into large cell aggregates, eventually detaching from the substrate after continued culture for around 5-6 weeks *in vitro* (Thiry et al., 2022; Lépine et al., 2024). Presuming that βII-spectrin gaps represent a cytoskeletal rearrangement in response to cellular stress that then leads to detachment, we treated 2-week-old MNs with agents known to induce acute cellular stress: arsenite (Ratti et al. 2020), L-glutamate (Lépine et al., 2024; Shi et al., 2018) and staurosporine (Zhang et al., 2013) (Suppl. Fig. 3A). None of these acute 1 h treatments induced loss of axons (Suppl. Fig. 3B). Among them, only staurosporine—a potent, cell-permeable, reversible, ATP-competitive and broad spectrum inhibitor of protein kinases (Nakano & Omura, 2009) significantly increased the percentage of axons with gaps (Fig. 3B and D). The increase in axons with βII-spec-gaps induced by 1 h of staurosporine treatment was maintained even 24 hrs and 72 hrs after treatment removal (Fig. 3C and Suppl. Fig. 3C and D), implying that 1 hour of staurosporine produces a persistent effect on the formation of βII-spec-gaps. It is important to note that no appreciable axonal loss was observed at these survival times (Suppl. Fig. 3D). Apart from their increase in number, βII-spectrin gaps and patches induced by 1 h staurosporine were indistinguishable from those present in vehicle controls (DMSO treated, Fig. 3E). Thanks to this, most subsequent analyses were conducted in staurosporine-treated MNs to ensure acute temporal control over the formation of these structures.

**Figure 3.**
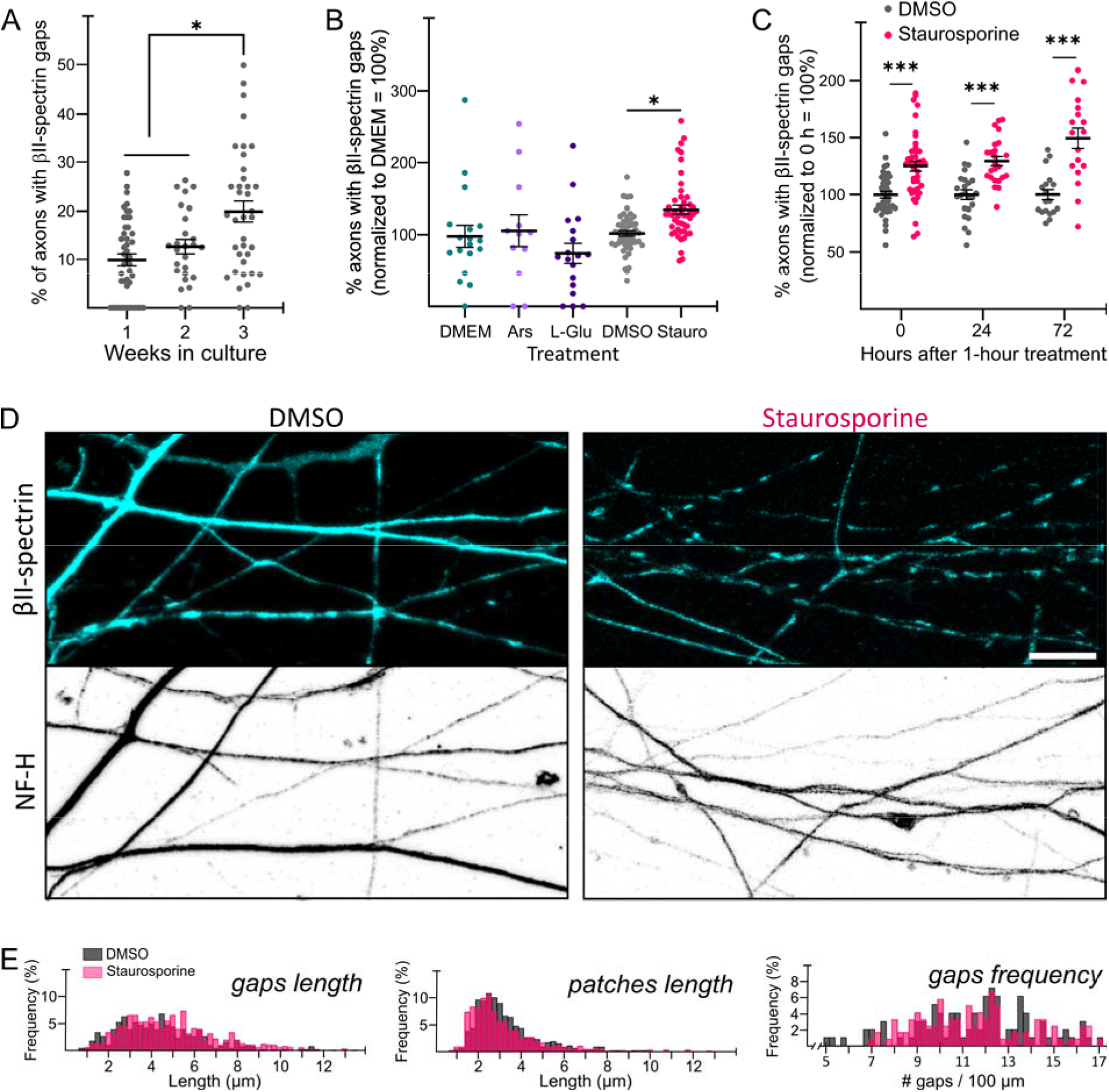
Staurosporine acutely induces βII-spectrin gaps-and-patches patterns. (**A**) Graph showing the percentage of axons with βII-spectrin gaps in iPSC-derived MNs cultured for 1, 2 and 3 weeks, and treated for 1 h with vehicle (DMSO). One way ANOVA, *:p < 0.05. (**B**) % of axons with βII-spectrin gaps (normalized to control) in 2-week-old MNs treated for 1 h with arsenite (2 μM), L-glutamate (0.05 mM) and staurosporine (0.1μM), or the control vehicles, and fixed immediately afterward. One way ANOVA, *:p < 0.05. (**C**) % of axons with βII-spectrin gaps (normalized to time 0) of 2-week-old MNs treated for 1 h with DMSO or staurosporine (0.1μM) and fixed immediately (0), or 24 and 72 hrs later. One way ANOVA, ***:p<0.001. (**D**) Confocal images of 2-week-old MNs treated for 1 h with DMSO or staurosporine (0.1μM) and immunostained for βII-spectrin and Neurofilament H. Scale bar = 10 μm. (**E**) Histograms showing percentual frequency of gaps length, patches length and gap frequency per 100 μm, comparing DMSO and staurosporine.

Given the clinical relevance of lower MN susceptibility in motor neuron diseases, we sought to determine whether ALS-related mutations influence the percentage of axons exhibiting βII-spectrin gaps. To this end, we differentiated three homozygous knock-in (KI) iPSC lines harboring mutations in FUS (H517Q), TARDBP/TDP43 (A382T) and SOD1 (G93A and D90A/G93A) (Deneault et al., 2022; Lépine et al., 2024) into iPSC-derived MNs and assessed them under basal and staurosporine treatment conditions. Interestingly, the proportion of axons with gaps was similar among the isogenic control (AIW002-02) and the ALS-related mutants under basal conditions. Likewise, staurosporine treatment produced a comparable increase in the proportion of axons with βII-spec-gaps across cell lines (Suppl. Fig. 3E). Under these conditions total axonal length remained unchanged (Suppl. Fig. 3F). These findings indicate that βII-spectrin distribution and its response to staurosporine are unaffected by the presence of ALS-related mutations.

### βII-spectrin gaps formation is not correlated with proteolytic cleavage of spectrins, nor with the activity of calpains or caspase-3

Spectrins can be cleaved by calpains and caspases and this processing is modulated during cell remodelling, such as during axon regeneration (Girouard et al., 2018). We hypothesized that regulated activity of these proteases could explain the staurosporine-induced gaps by locally targeting spectrins and disrupting the MPS lattice. Spectrin cleavage products are stable enough to appear as lower molecular weight bands by western blot. To perform this analysis, we treated 2-week-old MNs with vehicle or staurosporine for 1 h, and lysed them either immediately after washout, or 6 hrs later following a medium change (Suppl. Fig. 4A). First, we performed immunoblots analysis for αII- and βII-spectrin to determine whether spectrins are targeted by proteases during gap formation. Neither αII-nor βII-spectrin showed significant differences between conditions in the full length band or in the lower molecular weight bands expected as a result of proteolytic cleavage (Fig. 4A and Suppl. Fig. 4B and D). Thus, neither caspases nor calpains appear to target spectrins during staurosporine-induced gap formation.

**Figure 4.**
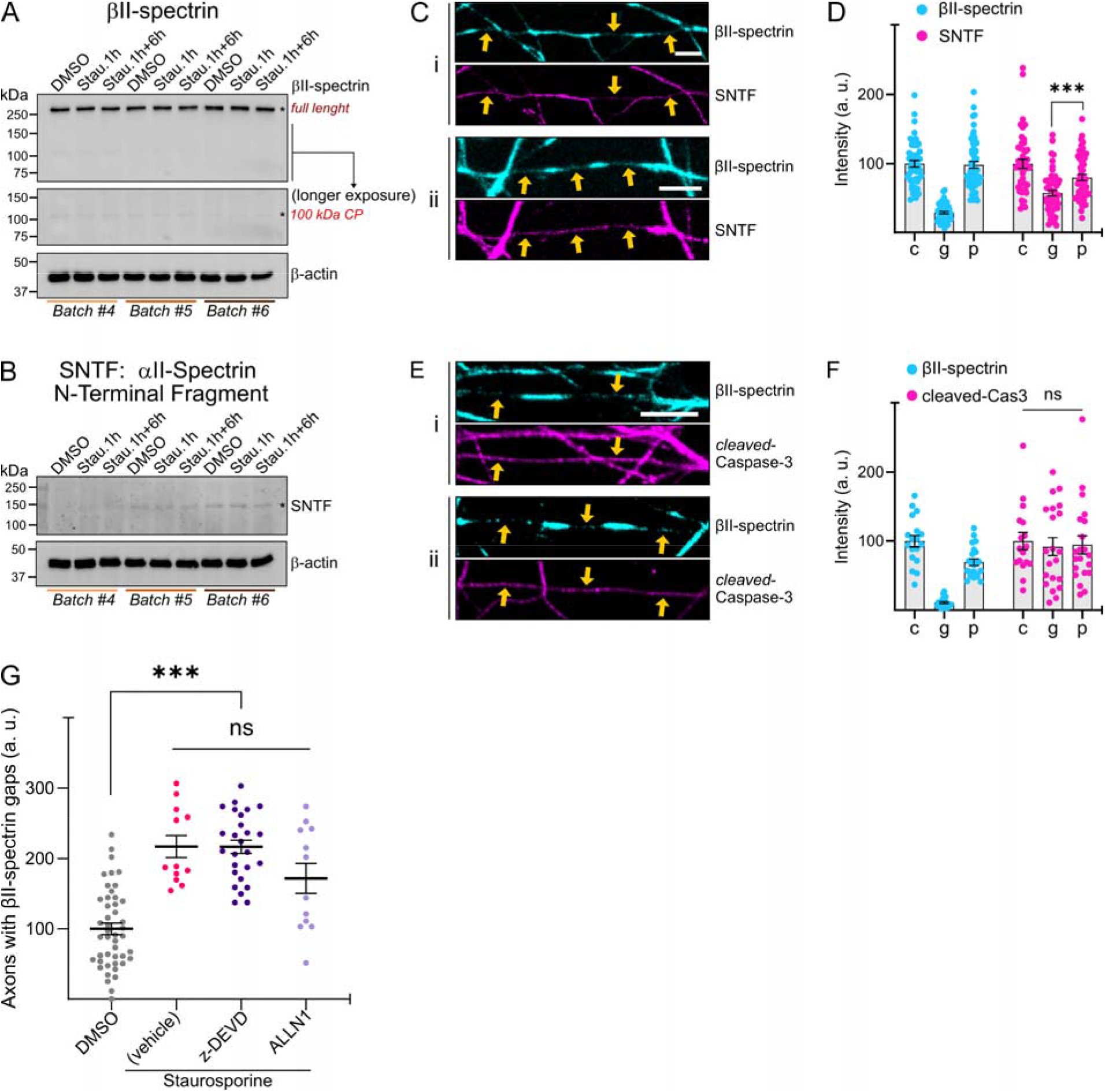
Proteolytic activity is not likely related to the formation of βII-spectrin gaps. (**A**) Western blot against βII-spectrin and β-actin in 2-week-old MN cultures from three independent differentiations (batches) treated with vehicle control (DMSO) or staurosporine for 1 h (stau. 1h), and then either pelleted immediately or 6 hrs later following a medium change (stau. 1h+6h). (**B**) Western blot against SNTF (αII-Spectrin N-Terminal Fragment) and β-actin in the same samples as in panel A. (**C-D**). Representative images (**C**) and quantification (**D**) of βII-spectrin and SNTF in continuous axons (c), in gaps (g) and patches (p) of staurosporine-treated 2-week-old MNs. Two way ANOVA, ns: not significant. (**E-F**). Representative images (**E**) and quantification (**F**) of βII-spectrin and cleaved-Caspase-3 in continuous axons (c), in gaps (g) and patches (p) of staurosporine-treated 2-week-old MNs. Two way ANOVA, ***: p<0.001. (**G**) Quantification of axons with βII-spectrin gaps in vehicle (DMSO) and staurosporine-treated 2-week-old MNs with or without the inhibitors for caspase-3 (z-DEVD-fmk) or calpains (ALLN1). One way ANOVA, ***: p<0.001.

We further examined these samples with an antibody that recognizes an αII-spectrin fragment produced by calpain cleavage. The SNTF antibody (Spectrin N-Terminal Fragment, (Roberts-Lewis et al., 1994)) targets a neoepitope of 6 amino acids at the N-terminus of the stable, calpain-derived αII-spectrin fragment of ~150 kDa. We found no consistent changes after 1 h of staurosporine treatment, and highly variable changes 6 hrs after (Fig. 4B and Suppl. Fig. 4C). Since subtle changes in microdomains can be masked in whole-culture homogenates, we performed quantitative confocal imaging of 1 h staurosporine-treated axons immunostained with anti-SNTF, and compared SNTF signal intensity across continuous axons, and within gaps and patches. Unexpectedly, the SNTF signal in gaps was ~25% lower than in neighboring patches, which is inconsistent with the hypothesis of a local SNTF production in gaps (Fig. 4C and D). We conclude that αII-spectrin cleavage -SNTF generation-does not appear to be a relevant feature of gap formation.

Caspase-3 is the main executioner caspase in axons, and its active form is generated by cleavage of pro-Caspase-3, a process that can also be followed by western blots. We used two antibodies to detect cleaved-Caspase-3 and found that it was present at very low levels across all treatment conditions. In two of the three batches, both antibodies detected a modest induction of cleaved-Caspase-3, but the levels remained well below what was expected for a global induction by staurosporine (Suppl. Fig. 4E and F).

To assess this in more detail, we performed confocal imaging of 1 h staurosporine-treated axons immunostained with anti-cleaved-Caspase-3, and compared its intensity in continuous axons, gaps and patches. We found no significant differences in cleaved-Caspase-3 signals among continuous axons, gaps and patches (Fig. 4E and F). We conclude that the generation of cleaved-Caspase-3 does not appear to be a relevant feature of gap formation.

To further assess whether protease activity is required for staurosporine-induced gap formation, we applied a set of validated protease inhibitors. When added 1 h before staurosporine, inhibition of Caspase-3 (z-DEVD-fmk) or calpains (ALLN1) activity failed to alter the induction of axons with gaps (Fig. 4G). Taken together, these results suggest that neither the acute, local activity of caspases or calpains nor the degradation of spectrins is necessary for the emergence of MPS gaps.

### The MPS is absent in βII-spectrin gaps, but it is well organized within patches

Confocal imaging allowed us to identify and quantify micrometer-sized gaps and patches in the βII-spectrin lattice within axons of MNs in bulk cultures. However, the resolution achievable through this approach precludes the study of structural features of the MPS. To overcome this, we next used STED nanoscopy to examine the nanoscale organization of βII-spectrin in different regions of these axons. First, we observed that βII-spectrin in axons of 2-week-old MNs exhibited the typical organization of the MPS, and surprisingly, that the MPS was also present in βII-spec-patches—both in vehicle- and staurosporine-treated MNs (Fig. 5A). The extent of MPS organization was assessed quantitatively by two methods: 1) the autocorrelation analyses of 2 μm segments, measuring the mean amplitude of the first two peaks of the autocorrelogram (as used elsewhere, (Zhong et al., 2014); and 2) the open-source image analysis tool “Gollum”, designed for the automated quantification of the quality of protein periodic structures, as detailed in the Materials and Methods section (Barabas et al. 2017). These measurements confirmed that the MPS is present in both continuous axons (cont.) and in βII-spec-patches (patch, Fig. 5B and C), although its organization in the patches tends to be less regular. Interestingly, a 1 h treatment with staurosporine increased the organization in continuous axons (cont., Fig. 5C).

**Figure 5.**
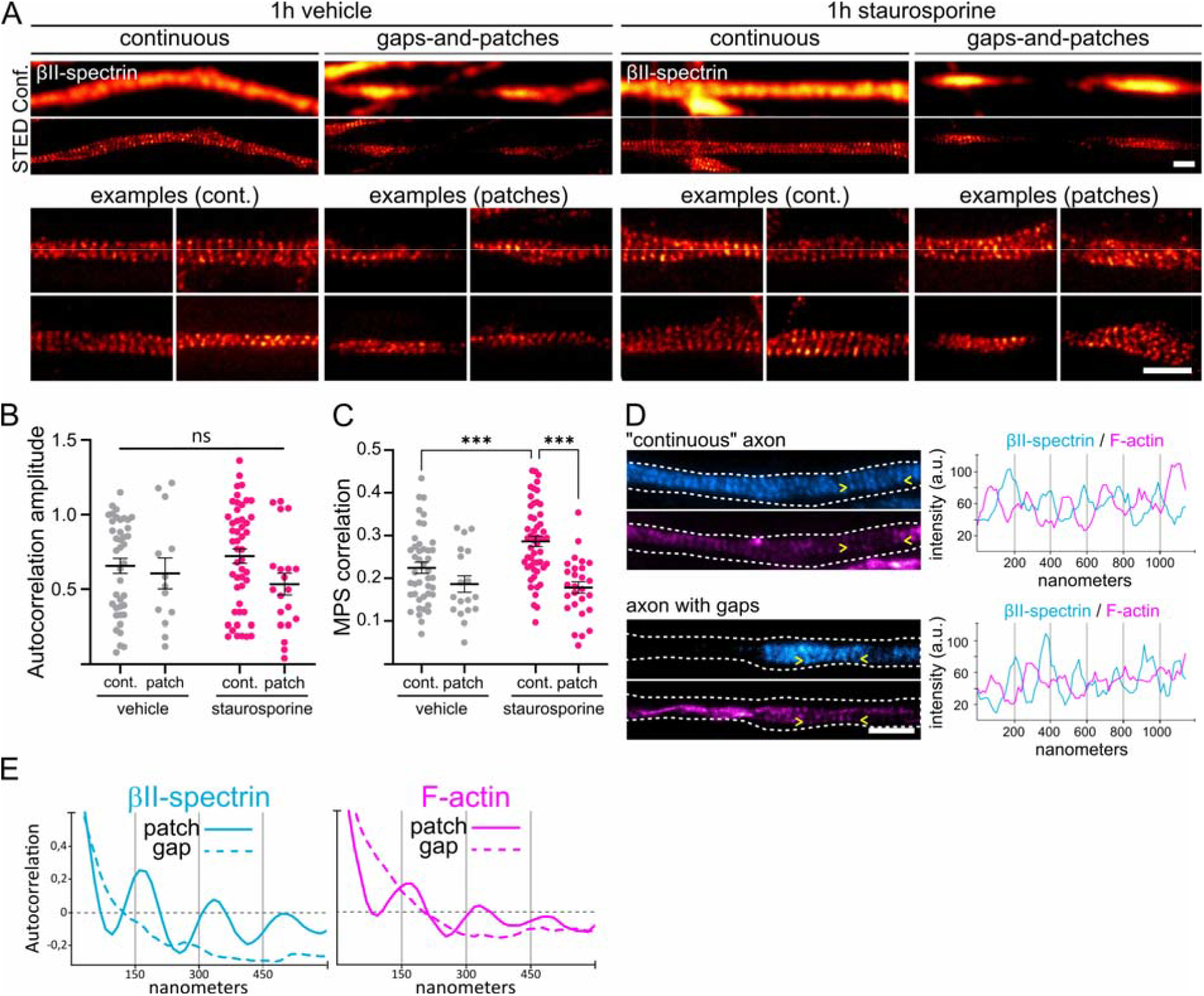
The MPS is absent in βII-spectrin gaps, but it is well organized within patches. (**A**) Representative images of βII-spectrin acquired using confocal and STED microscopy in axons with continuous βII-spectrin distribution and in axons with βII-spectrin gaps-and-patches pattern, under both control conditions and acute staurosporine treatment. Scale bars = 1 μm. (**B**) Autocorrelation amplitude of the intensity profiles along the different regions of interest and treatments. Mean ± SEM. ns: not significant. (**C**) MPS correlation analyses by Gollum within the different regions of interest and treatments. Mean ± SEM. Two way ANOVA, ***: p<0.001. (**D**) STED images of double staining for βII-spectrin and F-actin showed the expected anti-phase organization in both continuous axons and patches. Representative images (left) and corresponding intensity profiles between the arrowheads (right). Scale bar = 1 μm. (**E**) Autocorrelation analyses of βII-spectrin and F-actin in patches and gaps. The traces represent the mean autocorrelation from 17 pairs of patches and gaps.

Next, we investigated whether the absence of βII-spectrin in gaps reflects a loss of the underlying MPS, considering that βIII- and βIV-spectrin might replace βII-spectrin and hence allow MPS formation in its absence (Xu et al., 2013; Han et al., 2017). Since actin is a core component of the MPS regardless of the β-spectrin subunit composition, we evaluated whether actin filaments exhibited periodic organization within gaps as an indicator of MPS presence. STED imaging of phalloidin-stained MN axons revealed that actin is periodically organized in “continuous” axons and βII-spect-patches displaying periodic βII-spectrin, consistent with the presence of the MPS (Fig. 5D and E). In contrast, periodic F-actin organization was completely absent in the βII-spect-gaps, indicating that these microdomains lack an MPS (Fig. 5E). In support of the notion that βII-spec-gaps represent a lack of the MPS, αII-spectrin—another core component of the MPS and the only α-spectrin expressed in neurons—also displayed gaps and patches with a size and frequency distribution similar to those observed for βII-spectrin, suggesting their co-localization (Supp. Fig. 2A and C). Notably, αII-spectrin patches exhibited an MPS structure with a degree of organization comparable to that found in continuous axons (Supp. Fig. 5A).

We extended these super-recolution analyses to study the MPS on the (KI) iPSC lines harboring mutations in TARDBP/TDP43 (A382T) and SOD1 (D90A/G93A), in comparison to their isogenic control AIW002-02, focusing on axonal sections with a continuous βII-spectrin distribution. We found that between ~80% of regions had an MPS, with no difference among KI lines, and all showing a small increase when treated with staurosporine (Suppl. Fig. 5B). In MPS-bearing regions, the period found was ~190 nm, with no differences among cell lines or comparing DMSO- or staurosporine-treated (Suppl. Fig. 5C). Moreover, the extent of organization of the MPS did not show differences among these cell lines and the control line, and while the organization showed a small increase when treated with staurosporine in all lines, this change did not reach statistical significance (Suppl. Fig. 5D).

Taken together, STED nanoscopy enabled the characterization of MPS organization in human iPSC-derived MNs, demonstrating that βII-spectrin-patches contain a well-organized MPS, whereas βII-spect-gaps indeed lack an MPS. Also, ALS-related mutations TARDBP/TDP-43A382T and SOD1-D90A/G93A do not seem to affect MPS presence, period or organization.

### Gaps and patches preferentially occur in the middle section of axons

We next speculated that determining the proximal-to-distal location of the gap-and-patch pattern, together with a detailed analysis of protein signal intensities along individual axons, could provide insight into the formation of this pattern. We treated “density gradient” cultures with staurosporine for 1 h -to increase the abundance of βII-spectrin gaps- and fixed them immediately afterwards. Axons were imaged within 100 μm x 100 μm ROIs, and the number of gaps per section was quantified (Supp. Fig. 6A). Under these conditions, 72 % of sampled axons presented a gap-and-patch pattern. Among these, the majority (46 %) showed gaps exclusively in the medial section, while an additional 15 % displayed gaps in both the medial and distal sections (Fig. 6A). The graph in Fig. 6B shows the number of patches found in each section (normalized by length) for all 26 axons examined, where light grey traces and green dots highlight the axons that presented more gap-and-patches in the medial section. This analysis showed that the gap-and-patch pattern occurs preferentially in the medial section of 2-week-old MN axons. We then investigated whether βII-spectrin intensity along individual axons with a gap-and-patch pattern confined to their middle portion could provide insights into predicting the location of this pattern. To do this, we compared βII-spectrin intensity in gaps and patches with respect to flanking proximal and distal segments, which lacked gaps (Fig. 6C). Interestingly, we found that βII-spectrin intensity within patches was similar to that of the nearest proximal continuous section and higher than that of the nearest distal continuous section, whereas intensity within gaps was slightly lower than in the nearest distal continuous section (Fig. 6C and D). One possible explanation for the differential concentration of βII-spectrin observed above involves differences in the diffusion rates of “free” vs “MPS-bound” βII-spectrin, as recently demonstrated by *in vivo* FRAP experiments in *C. elegans* (Glomb et al., 2023). As patches have an organized MPS, it is reasonable to consider that these patches arise within previously unorganized sections of the axon. Since “free” βII-spectrin incorporated into the forming MPS would stop diffusing, each patch acts as a sink of βII-spectrin. This leads to increased βII-spectrin intensity within patches, and a corresponding depletion in the neighboring regions, which appear as gaps. The resulting gap-and-patch pattern may represent an intermediate stage in the assembly of a mature, long-range MPS lattice. This model accounts for most of the observations presented so far, namely the presence of an MPS in patches (Fig. 5A), its absence in gaps (Fig. 5D and E), and the microdomain changes in βII-spectrin intensity along axons with a gap-and-patch pattern (Fig. 6D).

**Figure 6.**
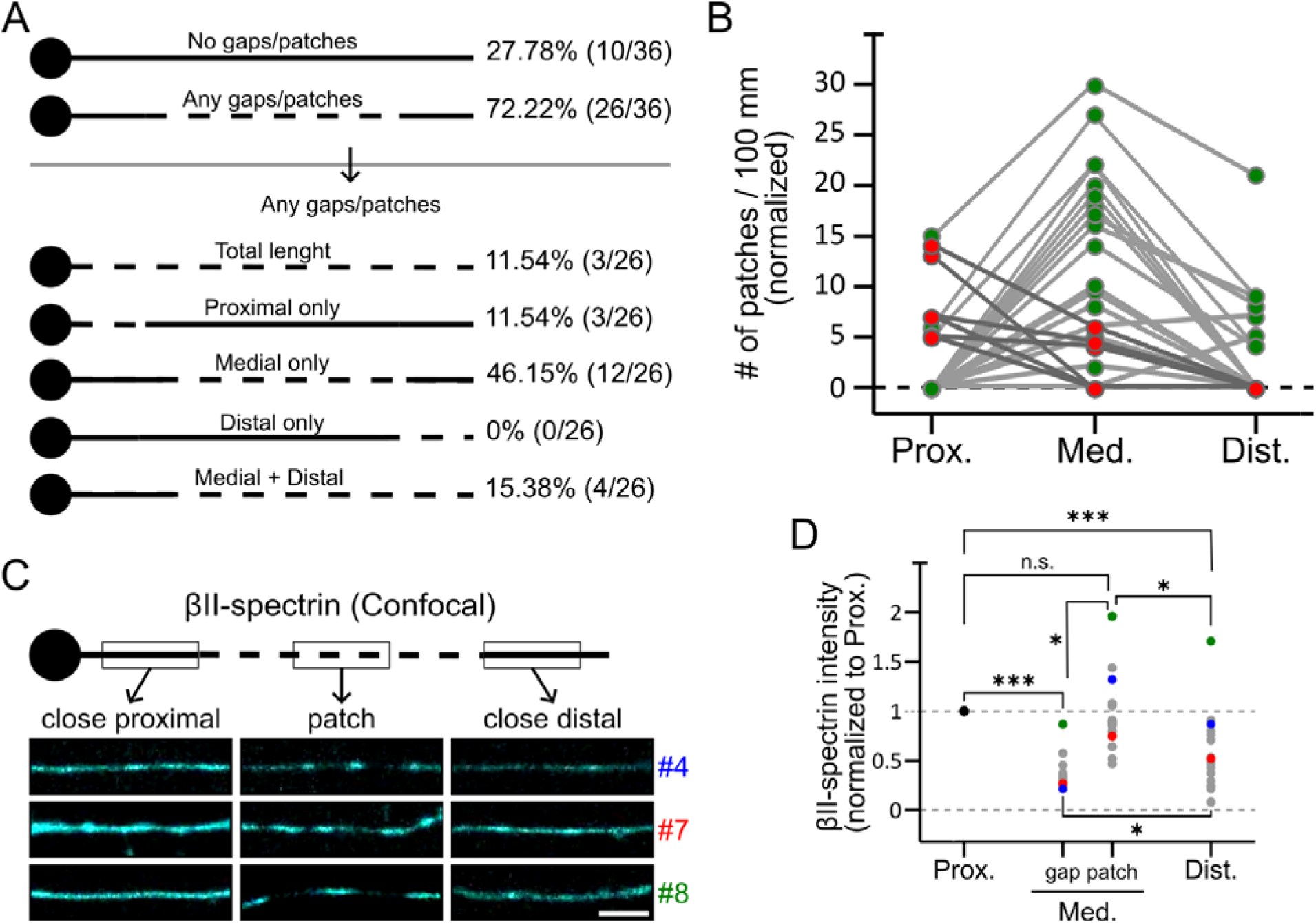
Gaps and patches preferentially occur in the medial portion of axons. (**A**) Schematic representation of the location of the gaps and patches pattern along individual axons (left) and its observed frequency (right). (**B**) Normalized number of patches per 100 μm in the proximal, medial and distal portions. Each trace represents an individual axon. Most axons showed more gaps and patches in the medial portion (light gray traces and green dots), while only a few showed more in the proximal region (dark gray traces and red dots). (**C**) Schematic (top) and representative confocal images of three βII-spectrin-stained axons with a gap-and-patch pattern confined to the medial sections, showing proximal, medial and distal sections (bottom). Scale bar = 1 μm. (**D**) Mean normalized βII-spectrin intensity in the proximal, medial (with gaps and patches pattern), and distal sections. Values were normalized to the proximal intensity of each axon. Colored dots correspond to the three examples shown in panel C. ns: not significant; *: p<0.05; ***: p<0.001.

In line with this conclusion, recent reports show the existence of nascent MPS segments, similar to our “patches”, in distal segments of hippocampal axons in culture (Hofmann et al., 2022; Boyer et al., 2026). In particular, Hofmann and colleagues found that calpain inhibition during axon regeneration did not alter the amount of patches in the new axonal segments, but they suffered a slight increase in their size. Taking this into account, we re-evaluated caspase and calpain inhibition experiments to determine whether protease inhibition altered the length of patches produced during 1 hour of staurosporine treatment. We found that staurosporine decreased by approximately ~10% de length of patches (which agrees with the comparative histograms for patch length shown in Fig. 3E), while DEVD and ALLN1 reversed this small change (Supp. Fig. 6B). These results can be interpreted in light of limited spectrin disponibility during MPS assembly (i.e. patch formation). On one hand, a limited amount of spectrin makes the extra patches induced during staurosporine treatment to be smaller. On the other hand, protease inhibitors revert that trend, by increasing the pool of spectrin availability.

### Differential MPS organization along axons suggests that patches represent nascent MPS assemblages

To validate the hypothesis that the gap-and-patch pattern represent an intermediate stage in the assembly of a mature, long-range MPS lattice, we asked whether the periodical organization of βII-spectrin in the MPS is maintained across the proximal, medial, and distal sections of individual axons exhibiting a gap-and-patch pattern in the medial segment (schematics in Fig. 7A). Specifically, we compared *MPS correlation coefficients* calculated on STED images from the continuous segment in the proximal portion, patches in the middle portion, and the continuous segment in the distal portion along individual axons (Fig. 7A). As observed in bulk MN cultures, the patches exhibit a MPS organization comparable to that of the continuous segment in the proximal region (Fig. 7A and B). In contrast, the distal portion adjacent to the gap-and-patch pattern lacked an organized MPS (Fig. 7A and B). We also noted that patch borders were not sharply defined. Instead, they presented incomplete transverse βII-spectrin labeling while maintaining a longitudinal periodic distribution characteristic of the MPS (Fig. 7C and D). Notably, gaps with higher βII-spectrin intensity under confocal imaging showed increased autocorrelation amplitude of this longitudinal periodic arrangement (Fig. 7D). Since axons grow by adding membrane at their distal tips (Pfenninger et al., 2003), distal sections represent more recently formed axonal compartments. Taken together, these observations suggest that patches in the medial section represent *de novo* MPS assembly within previously unorganized, and newer, axonal regions.

**Figure 7.**
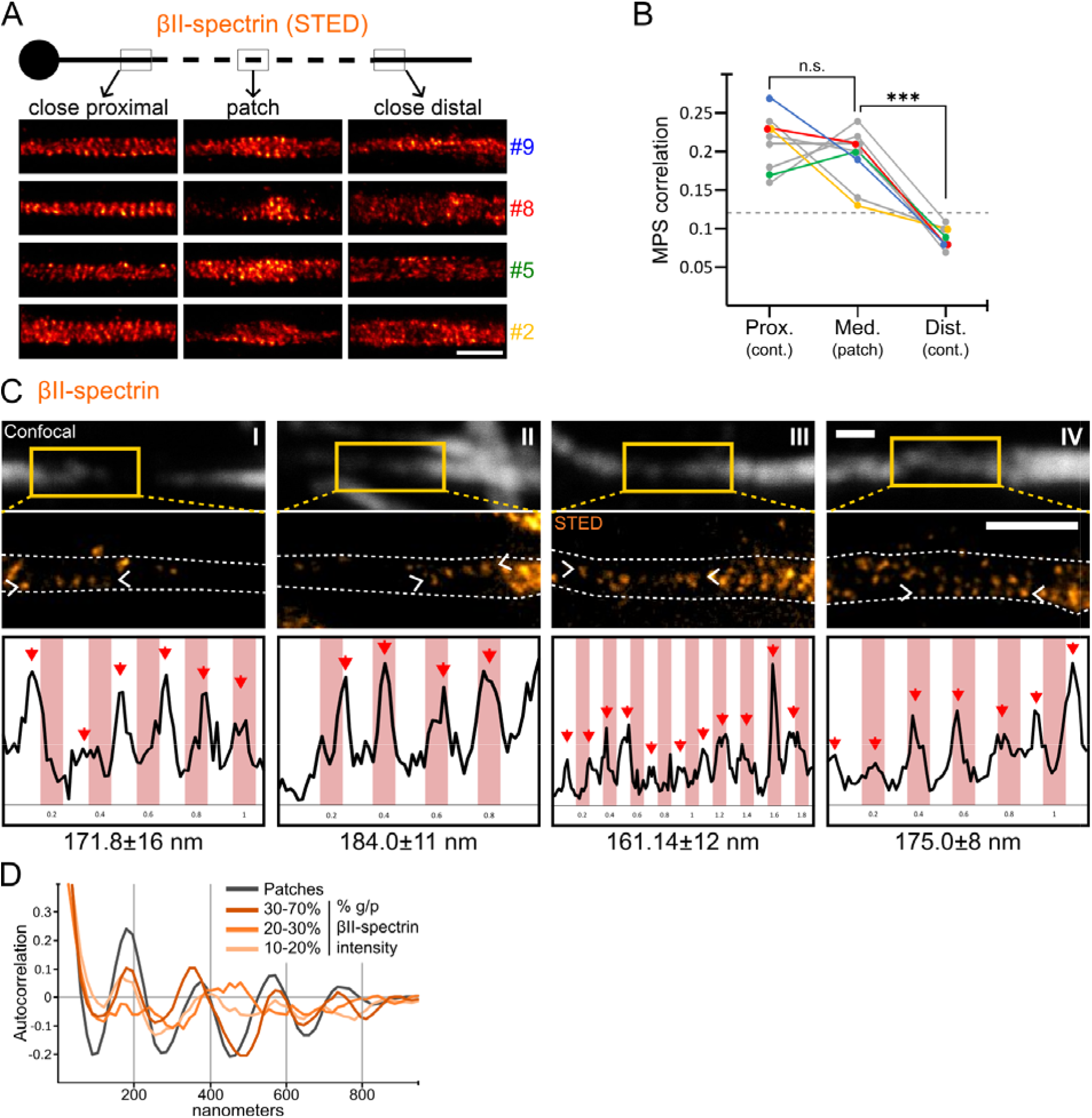
Patches may represent nascent MPS assemblages. (**A**) Representative STED images of four individual axons immunostained against βII-spectrin, showing continuous segments in the proximal and distal portions, and patch segments in the medial portions. Scale bar = 1 μm. (**B)** MPS correlation values of 12 individual axons in the continuous segments of the proximal and distal portions, and patch segments in the medial portion. Colored traces correspond to the axons shown in panel A. The dashed line at 0.12 indicates the threshold above which a periodic distribution is visually discernible. One way ANOVA, ns: not significant; ***: p<0.001. (**C**) Representative confocal and STED images of four individual gap-patch borders immunostained for βII-spectrin. White arrowheads indicate the locations of the intensity profiles. Red arrows marked the regions used to calculate the mean period for each section, as shown below. Pink stripes in the background are separated by 200 nm and were drawn for visual reference. Scale bars = 1 μm. (**D**) Mean autocorrelation curves of longitudinal βII-spectrin structures across sample groups with varying βII-spectrin intensity in the gaps, expressed as a percentage of the intensity found in the neighboring patches. A mean AC curve of patches (grey) is shown for reference.

### Acute pharmacological sequestration of actin monomers prevents the formation of staurosporine-induced βII-spectrin patches

It has been suggested that the nucleation of new actin filaments is important for MPS formation—rather than elongation of existing filaments (Qu et al., 2017). We speculated that staurosporine may induce the gap-and-patch pattern by stimulating the formation of new actin filaments. To acutely interfere with actin polymerization during staurosporine treatment, we used Latrunculin A (LatA, 1 μM), a small molecule known to bind actin monomers and prevent their polymerization into new filaments or incorporation into existing ones (Morton et al., 2000). LatA was added 1 h before staurosporine and remained present during the 1-hour staurosporine treatment; cultures were then immediately fixed and immunostained. LatA prevented the staurosporine-induced formation of gaps, without altering the baseline occurrence of gaps in the absence of staurosporine (Fig. 8A and B). Quantification of global axonal F-actin staining in these cultures confirmed that treatment with LatA reduced the levels of actin filaments (Fig. 8C). On the other hand, staurosporine alone significantly increased the levels of F-actin staining, further supporting the hypothesis that staurosporine stimulates the formation of new actin filaments (Fig. 8C). Interestingly, while LatA treatment blocked the formation of staurosporine-induced patches, it did not alter MPS organization in continuous axons with a similar intensity of βII-spectrin or in the preserved patches -likely present before LatA-(Fig. 8D and E). These results indicate that, at the dose used, LatA does not disrupt existing MPS structures, but only the generation of new MPS segments.

**Figure 8.**
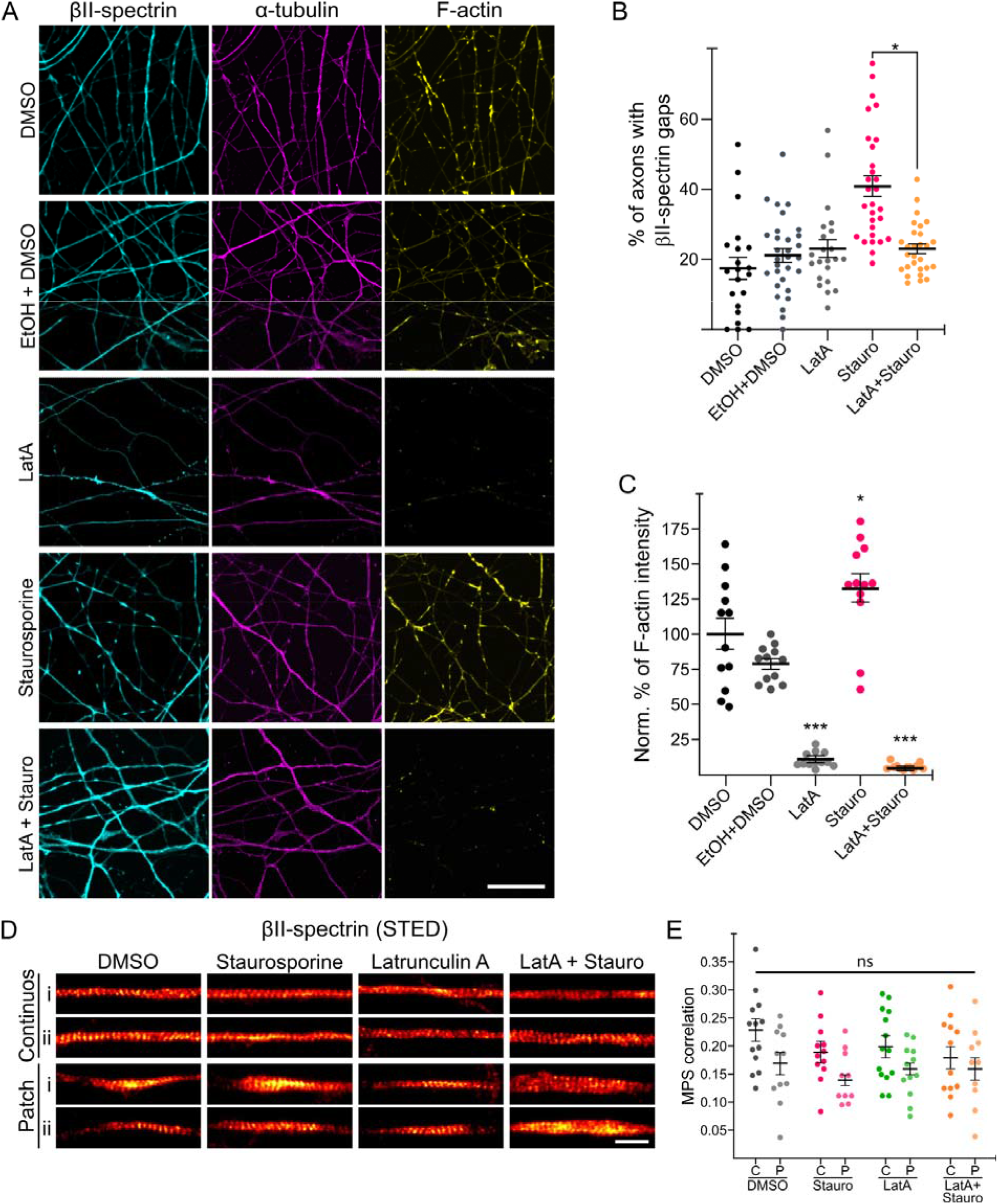
Latrunculin A prevents staurosporine-induced gaps-and-patches pattern formation. (**A**) Representative confocal images of 2-week-old MNs treated with vehicle (DMSO or EtOH+DMSO), latrunculin A (LatA), staurosporine (Stauro), or latrunculin A + staurosporine (LatA+Stauro). Cells were immunostained for βII-spectrin, α-tubulin and F-actin (phalloidin). Scale bar = 20 μm. (**B**) Percentage of axons with βII-spectrin gaps in 2-week-old MNs treated groups shown in panel (A). Mean ± SEM. *: p<0.05. Tukey post hoc test. (**C**) Normalized percentage of F-actin intensity in 2-week-old MNs treated groups shown in panel (A). Mean ± SEM. *: p<0.05, ***: p<0.001. Tukey post hoc test. (**D**) Representative STED images of MNs treated with vehicle (DMSO), staurosporine alone (Stauro), latrunculin A alone (LatA) or latrunculin-A + staurosporine (LatA+Stauro), and immunostained for βII-spectrin. Two examples are shown for continuous axonal sections (top rows) and for patches (bottom rows). Scale bar = 1 μm. (**E**) MPS correlation analyses by Gollum within the different regions of interest and treatments shown in panel D (C: continuous, P: patches). ns: not significant. Mean ± SEM.

## Discussion

In this study, we provide a detailed characterization on the spatial organization of βII-spectrin and the membrane-associated periodic skeleton (MPS) in human iPSC-derived MNs. Using confocal imaging, we identified the existence of sharply demarcated interruptions in the otherwise continuous βII-spectrin lattice, forming a distinctive gap-and-patch pattern along axons. These interruptions, or “gaps,” which also lack αII-spectrin, occur interspersed with patches where the MPS is well preserved. The formation of βII-spectrin gaps increases with culture duration and can also be acutely induced by the kinase inhibitor staurosporine. Importantly, we showed that the gap-and-patch pattern is not linked to axonal degeneration or cytoskeletal disassembly but may instead reflect a dynamic and spatially regulated process of MPS formation and maturation. Additionally, we demonstrate that pharmacological inhibition of actin polymerization blocks gap formation, and further propose that actin nucleation is required for the initial MPS assembly.

To our knowledge, this is the first detailed study to map the distribution and organization of βII-spectrin and the MPS along human MN axons at nanoscale resolution — the existence of the MPS in human iPSC-derived MNs was briefly described previously (He et al., 2016). MN axons are the target of dysfunction and degeneration in various neurological conditions and neurodegenerative diseases. Because active remodeling of the MPS has been described in degenerating axons (Unsain, Bordenave, et al., 2018; G. Wang et al., 2019) and various ALS-related genes directly affect cytoskeletal dynamics (Castellanos-Montiel et al., 2020), we expected that a better understanding of the actin/spectrin cytoskeleton in motor neurons axons could provide clues regarding their vulnerability. Notably, we observed that ALS-related mutations in SOD1, TARDBP/TDP43 and FUS do not significantly affect either the baseline occurrence of βII-spectrin gaps or their induction by staurosporine, suggesting that the normal assembly process of the MPS is preserved in these disease models. In previous reports we have shown that human iPSC-derived MNs bearing ALS-related mutations exhibit certain disease phenotypes only when maintained in culture for extended periods (Deneault et al., 2022; Lépine et al., 2024), raising the question of how the MPS behaves in neurons cultured for longer periods. A more detailed analysis to determine whether MPS stability is altered in disease-relevant contexts should be examined in the future.

Initially, we speculated that the appearance of gaps could result from MPS disassembly via localized proteolytic degradation of its key components. Caspases and calpains are known to cleave spectrins during neuronal injury and axon degeneration (Zhang et al., 2009; Kobeissy et al., 2015), as well as during normal cell remodeling (Lee et al., 2001; Glantz et al., 2007; Girouard et al., 2018). However, our western blot and immunofluorescence analyses revealed no consistent increase in spectrin cleavage products, nor in markers of caspase or calpain activity during staurosporine-induced gap formation. Moreover, the use of pharmacological inhibitors of these proteases failed to prevent the formation of gaps-and-patches patterns. These results argue against the involvement of protease-mediated spectrin degradation that could explain the formation of gaps. On the contrary, our data implies that βII-spectrin patches represent nascent MPS assemblies arising from previously unstructured spectrin-actin complexes along newer axonal domains. In line with this hypothesis, others have recently described the appearance of short MPS segments in medial and distal portions of hippocampal axons in culture (Hofmann et al., 2022; Boyer et al., 2026). On the possible role of proteases in such structures, Hofmann and colleagues found that acute treatment with calpain inhibitors right before axotomy lead to an increase in percentage of periodic spectrin (referred by authors as “periodicity”) in the regenerated axons in a 2-hour period. Interestingly, the spectrin patches did not increase in number, but they increased in size. This indicates that in the particular situation of axonal regeneration, calpain activity puts a brake into MPS formation within patches. When we thus examined staurosporine-patches in the presence of caspase and calpain inhibitors, we found that these prevented the slight drop in size produced by staurosporine. We conclude that indeed protease inhibitors have an impact on patch dynamics, likely by changing the availability of spectrins. However, in our case this effect was considerably smaller than the one observed in regenerated axons (changes of ~10% vs ~100%, respectively).

Calpain and caspase activity has been shown to be relevant in other aspects of MPS biology (Heller et al., 2025; Fei et al., 2026). Heller and colleagues found that calpain activity contributes to the steady-state dynamics of spectrin exchange in a mature MPS lattice. More recently, Fei and colleagues describe a relevant role for calpains whenever massive endocytosis (of any kind) is engaged experimentally. Interestingly, all these studies, including ours, examined calpains roles in MPS in different scenarios. Altogether, these results are not contradictory among them, and provide valuable complementary information about the roles of calpains (and caspases) in MPS assembly, growth, maintenance and remodeling.

Our detailed super-resolution analyses further support the notion that the patches represent nascent MPS stretches. In axons displaying a medial gap-and-patch pattern, MPS organization was robust in both the proximal region and in the patches, but absent in the distal axon, Interestingly, the borders of patches often displayed an incomplete transverse coverage of βII-spectrin while maintaining longitudinal periodicity, suggestive of a transitional assembly zone. In support of this model, we observed that patches had βII-spectrin levels similar to proximal continuous regions. In turn, accumulation of spectrin within patches depletes surrounding regions and as a result gaps are evidenced between patches: transported spectrin could be locally sequestered into the nascent MPS in patches, which halts its diffusion and effectively depletes the surrounding axonal domains -and gaps emerge. This is consistent with recent *in vivo* data in *C. elegans* and hippocampal neurons in culture showing that spectrin diffusion is halted upon incorporation into the MPS (Glomb et al., 2023; Boyer et al., 2026). This dynamic mechanistic model can explain the appearance of a long gap-and-patch pattern found in relatively new portions of the axon. The validity of such a model could be further supported by live-imaging approaches with a tight control of expression levels of tagged βII-spectrin, as was recently developed in the mouse (Heller et al., 2025).

Previous work in rodent hippocampal neurons has described the progressive maturation of the MPS in developing axons (Hofmann et al., 2022; Zhong et al., 2014), and the mechanistic details of its initial assembly are beginning to be elucidated (Bodas et al., 2025; Boyer et al., 2026). All these studies, including the present one, agree on the initial observation by Zhong and colleagues that extending axons in culture have a better organized MPS proximal to the cell soma, and decrease gradually in moving towards the distal portions. Upon this observation, authors initially implied that new MPS segments were simply being added to the existing MPS, in a proximal-to-distal fashion (Hofmann et al., 2022; Zhong et al., 2014). However, this mechanism of MPS growth was later challenged by the observation that in distal regions devoid of a continuous MPS, βII-spectrin accumulations, called “patches”, represent immature MPS structures (Hofmann et al., 2022). Given these “patches” increase in size and periodical organization in a distal-to-proximal fashion, it had been proposed that new MPS segments would coalesce with older segments, giving rise to a continuous MPS. The “patches” described by Hofmann and colleagues in hippocampal neurons differ from the ones found by us in motor neurons in at least two major aspects. First, patches reported by us, once noticeable by confocal microscopy, have a complete MPS organization (i.e. periodic βII-spectrin occupies the complete width of the axon), whereas those described by Hofmann and colleagues do not cover the entire width and resemble the structures we found in the gap-patch borders. Second, the patches described by us in motor neurons arise in a gap-and-patch pattern, whereas patches shown by Hofmann and colleagues seemed to be isolated features. It is possible that axon extension rate and MPS components synthesis and trafficking rates can explain the differences observed in these cell types. However, it is worth mentioning that the observations by Hofmann and colleagues were performed in methanol-fixed cells, whereas ours were fixed in paraformaldehyde-sucrose in PBS. It is possible that some membrane-bound cytoskeletal assemblages are more noticeable in methanol fixation since it removes “unbound” components.

While this manuscript was in preparation, a preprint was published showing a similar gap-and-patch pattern as the ones found by us, but in hippocampal cells (Boyer et al., 2026). However, this report suggests that the patches have no MPS structure, but rather an amorphous condensate of spectrins. On the other hand, we suggest that all three studies have found amorphous, spectrin rich structures that may serve as a reservoir of spectrin tetramers that later diffuse and incorporate into the MPS. These structures that we suggest to have equivalent functions are: the accumulation of spectrin found in the growth-cone center (Hofmann et al., 2022), the “patches” in the axonal shaft (Boyer et al., 2026) and axonal enlargements shown here (Fig. 1). Further experiments would be needed to confirm this proposed function and equivalence of these structures. Taken together, in spite of the above mentioned differences, all three studies agree that the MPS grow by the maturation of “patches” of MPS/spectrin that then coalesce with the existent, continuous proximal MPS by lateral growth.

We further showed that preventing actin polymerization by sequestering G-actin with LatA prevents the formation of staurosporine-induced βII-spectrin patches without affecting the structure of pre-existing MPS regions. This indicates that the formation of the gap-and-patch pattern requires the assembly of actin filaments. The importance of actin dynamics for MPS stability was suggested many times using pharmacological interventions (Zhong et al., 2014; Leterrier et al., 2015; Unsain, Bordenave, et al., 2018; Abouelezz et al., 2019) and work in the fruit fly incorporated genetic approaches to highlight the particular importance of actin nucleation (Qu et al., 2017). While this manuscript was in preparation, a preprint was published in sensory neurons in culture supporting the notion that actin nucleation is necessary for MPS initiation (Bodas et al., 2025). These studies and our own results (Fig. 8), have shown that the actin filaments in the MPS are particularly resistant to drugs that alter filament extension, suggesting that their turnover is relatively low compared to other F-actin structures present in the axon. Most F-actin is lost upon sequestration of G-actin for 2 hours (by LatA) indicative of their high turnover -as a filament continues to lose monomers from the pointed-end but fails to incorporate monomers at the barbed-end, the filament get shorter, and can eventually disappear. The time frame of this process is indicative of its turnover rate. On the other hand, 2 hours of LatA did not alter the MPS, indicating that F-Actin in the MPS surfers negligible or none turnover during the 2-hours period. Taken together, these reports and our findings suggest that actin filaments within the MPS are stable, and that new MPS segments require the polymerization of new actin filaments.

A striking observation in our study is the induction of βII-spectrin gaps by the broad-spectrum kinase inhibitor staurosporine. Apart from the *number*, and a small decrease in patch length, we could not find any other distinction between gaps-and-patches found in untreated cells and in cells treated acutely with staurosporine, so we have assumed that both features are essentially the same objects of study. However, as a caution note for any conclusion driven from negative data, we cannot rule out the possibility that these objects are essentially different from one another. In the meantime, we believe that the treatment with staurosporine is very useful experimentally because it allows for a tight temporal control of gaps-and-patches formation. The effect of staurosporine is robust and persists for at least 72 hours, suggesting the engagement of a lasting remodeling process rather than a transient perturbation. Staurosporine has been previously shown to promote neurite outgrowth, actin reorganization, and lamellipodial activity in several neuronal systems (Rasouly et al., 1992; Jalava et al., 1993; Sano et al., 1994; Kohno et al., 2015). Although initially described as a PKC inhibitor, most of these effects seem independent of this master kinase. Based on our findings that Latrunculin A (LatA) can prevent staurosporine-induced gaps, we hypothesize that staurosporine may directly or indirectly promote actin filament nucleation. One potential mechanism is the inhibition of LIM kinase 1 (LIMK1), a kinase that phosphorylates and inactivates ADF/cofilin. Dephosphorylated cofilin enhances actin filament turnover and can paradoxically increase total F-actin by generating new barbed ends for filament elongation (Mannherz et al., 2006). Therefore, staurosporine-induced cofilin activation could drive the formation of nascent MPS (patches) by stimulating the extension of new actin filaments. Furthermore, phosphorylation of βII-spectrin and adducin has been implicated in weakening the spectrin-actin network (Manno et al., 1995; Matsuoka et al., 1998; Bignone et al., 2007), hence it is possible that staurosporine could indirectly stabilize nascent MPS structures by direct inhibition of the kinases responsible for the phosphorylation of βII-spectrin and adducin. Finally, while this work was under revision, Heller and colleagues showed that inhibition of PKC stabilized actin filaments in rings and the MPS as a whole, which is consistent with our findings using staurosporine (Heller et al., 2025). Finding the kinase cascade(s) targeted during the staurosporine-induced gap-and-patch pattern in human motor neurons could be addressed in subsequent studies.

In summary, we report the discovery of a patterned MPS initiation in human motor neuron axons, characterized by alternating βII-spectrin gaps and patches. This arrangement increases as a function of culture time and is accelerated by staurosporine in an actin-dependent manner. Our results suggest that patches represent sites of *de novo* MPS assembly rather than remnants of a degraded lattice. These findings provide new insights into the spatial and temporal regulation of the axonal cytoskeleton and lay the groundwork for future studies into how cytoskeletal dynamics is regulated and might be altered in neurodegenerative diseases such as ALS.

## Materials and Methods

### Human iPSCs lines

For this study we used the previously validated control iPSC line AIW002-02, reprogrammed from a 37-year-old Caucasian male (Chen et al., 2021). iPSCs were cultured on Matrigel (Corning, USA)-coated 100 mm dishes (Corning, USA) in mTeSR1 medium (STEMCELL Technologies, Canada) and passaged at ~80% confluence using Gentle Cell Dissociation Reagent (GCDR; STEMCELL Technologies, Canada). Detailed protocols for cell culture, characterization of pluripotency markers, qPCR-based assessment of chromosomal copy number, karyotyping, and virology testing are available in (Chen et al., 2021). Prior to starting the experiments, iPSCs were tested weekly and confirmed to be mycoplasma-free.

### Differentiation of iPSCs into motor neurons

AIW002-02 iPSCs were differentiated into motor neuron progenitor cells (MNPCs) as previously described (Deneault et al., 2022; Castellanos-Montiel et al., 2023; Lépine et al., 2024). After six days in MNPC Expansion medium (D18), cells were dissociated using GCDR and replated in MN Induction Medium containing 100 μM L-ascorbic acid (AA; Sigma-Aldrich, USA), 0.5 μM retinoic acid (RA; Sigma-Aldrich, USA), 0.1 μM purmorphamine (Pur; Sigma-Aldrich, USA), and 10 μM Y-27632 dihydrochloride (ROCK inhibitor; (STEMCELL Technologies, Canada). After 24 hrs, ROCK inhibitor was removed by performing a full medium change with fresh MN Induction Medium, which was subsequently replaced every other day. After six days in MN Induction Medium (D21), cells were dissociated into a single-cell suspension using Accutase (Thermo Fisher Scientific, USA) and replated in the required culture vessel for each experiment, with MN Induction and Maturation Medium containing 100 μM AA, 0.5 μM RA, 0.1 μM Pur, 0.1 μM Compound E (STEMCELL Technologies, Canada), 10 ng/mL brain-derived neurotrophic factor (BDNF; PeproTech, USA), 10 ng/mL ciliary neurotrophic factor (CNTF; PeproTech, USA), 10 ng/mL insulin-like growth factor (IGF-1; Peprotech, USA), and 10 μM ROCK inhibitor. After 24 hrs, ROCK inhibitor was removed by performing a full medium change with fresh MN Induction and Maturation Medium, which was subsequently replaced once a week. MNs were used for final experiments after two weeks in MN Induction and Maturation Medium, referred to throughout the study as 2 weeks-old MNs. For all experiments, three independent differentiations (“batches”) were performed using the iPSC lines for each of the four genotypes. All media were prepared in 1:1 Neurobasal:DMEM/F-12 (Thermo Fisher Scientific, USA) supplemented with 1X Antibiotic-Antimycotic (Thermo Fisher Scientific, USA), 0.5X N-2 (Thermo Fisher Scientific, USA), 0.5X B-27 (Thermofisher Scientific, USA) and 0.5X GlutaMAX (Thermo Fisher Scientific, USA).

### Coverslips coating, seeding and chemical treatment of neuronal cultures

A glass coverslip (#1, Karl Hecht Assistent, Germany) was placed in each well of a 24-well plate (Corning, USA). Each coverslip was then treated with 0.5 mL of 1X PBS (Wisent, Canada) containing 10 μg/mL poly-L-ornithine (PLO; Sigma-Aldrich, USA). After 24 hrs of PLO treatment, the coverslips were washed three times with 1X PBS, followed by an overnight treatment with 0.5 mL of DMEM/F-12 containing 5 μg/mL laminin (Sigma-Aldrich, USA). A total of 40,000 iPSC-derived MNPCs per well were plated and cultured for fourteen days in final differentiation medium, after which neurons were treated with the following reagents (more details as Supplementary information): Staurosporine (0.1 μM, Tocris Bioscience), L-glutamate (0.5 mM, Sigma-Aldrich), Sodium arsenite (0.25 μM, Sigma-Aldrich), Latrunculin A (1 μM, Cayman Chemical), ALLN-1 (5 μM, Cayman Chemical), Caspase-3 inhibitor(zDEVD-fmk, 50 μM, R&D Systems).

### Immunofluorescence staining

Samples were fixed for 20 min at room temperature (RT) in a fixation solution containing 4% formaldehyde (Thermo Fisher Scientific, USA) and 4% D-sucrose (Thermo Fisher Scientific, USA) in 1X PBS. They were then permeabilized with 0.2% Triton X-100 (Bioshop, Canada) in PBS for 5 min and blocked with two consecutive 5-min incubations in 0.1% Tween-20 (Bioshop, Canada) in PBS. Primary antibodies were incubated overnight at 4 °C, followed by a 2 hrs incubation with secondary antibodies at RT. When used, Phalloidin (Phalloidin-Atto 647N, Sigma-Aldrich) was co-incubated with the secondary antibodies. The primary antibodies used in this study were (more details as Supplementary information): anti-βII-spectrin (BD Biosciences), anti-α-II-spectrin and anti-βIII-tubulin (BioLegend), anti-αII-tubulin (Sigma-Aldrich), anti-Cleaved-Caspase-3 (Cell Signaling), anti-βIII-tubulin, anti-4.1N and anti-NF-H (Abcam), anti-SNTF and anti-NF-M(Millipore). And secondary antibodies were: anti-Goat anti-mouse IgG-STAR ORANGE, anti-Goat anti-rabbit IgG-STAR ORANGE and anti-Goat anti-rabbit IgG-STAR RED (Abberior Instruments GmbH); anti-Goat anti-mouse IgG-Atto 647N (Millipore), anti-Donkey anti-chicken IgY-Alexa488 (Jackson Immunoresearch), anti-Goat anti-rat IgG-Dylight488 (Abcam).

### Confocal microscopy and image analysis

Microscopy images were acquired at Centro de Micro y Nanoscopía de Córdoba (CEMINCO, Universidad Nacional de Córdoba, Argentina, https://ceminco.conicet.unc.edu.ar/) and at the Neuro Microscopy Core Facility (NMCF, McGill University, Canada). Confocal images were acquired using a Zeiss LSM 880 (NMCF) or a Zeiss LSM800 (CEMINCO) inverted microscope (AxioObserver platforms) equipped with a Plan-Apochromat 63x/1.40 NA oil immersion objective. Images were captured using internal photomultiplier tube (PMT) detectors and acquired through the ZEN software (version 14.0.27.201). Alexa Fluor 488, 568, and ATTO 647 dyes were sequentially excited using 488, 561, and 633 nm lasers, respectively. Emission was detected with spectral PMTs over ranges of 493–598 nm, 568–682 nm, and 638–759 nm, respectively. Pinhole diameter was set at 1.0 Airy Units. Z-stacks were acquired with an optimal step size yielding a final voxel size of 0.13 × 0.13 × 0.3 μm. Image averaging (2×) was applied during acquisition. Imaging conditions were set to avoid saturated pixels and allow observing linear changes in intensities. All imaging parameters were kept constant across conditions. Images were processed and further analysed (as indicated below) using FIJI software (Schindelin et al., 2012), with brightness and contrast adjusted uniformly only for visualization; no further processing was applied.

*Analysis of intensity*, the channel with βII-spectrin was used as reference to define the following regions of interest (ROIs) by drawing areas on: patches, gaps or the axons with continuous signal of this protein. In the latter, the sizes of areas covered about 3 μm in axon length, so as to be comparable to measures of gaps and patches. Background signal was thresholded to NaN. After that, the Mean Grey Value was measured for every channel. Mean grey values were normalized across experiments using the mean values found in ROIs belonging to the continuous signal group in each condition.

*Analysis of patch length, gap length, and gap frequency*, only the βII-spectrin channel was used as a reference. A segmented line was drawn along the axon; starting from a patch, each segment was measured as well as the total length of the process. These lengths were calculated using the x and y coordinates and the Pythagorean theorem. The frequency of gaps every 100 μm was then calculated.

#### Analysis of percentage of axons with βII-spectrin gaps

Whole frame z-stacks images taken with the 63X objective (101.21 μm x 101.21 μm) were first maximum-intensity projected. Then axons were counted as “with gaps” or “without gaps” in the signal of βII-spectrin if in a single axon, at least 3 consecutive gaps were found, forming a pattern. After that, percentages were calculated per image and normalized with the values of the control condition in a given experiment (DMSO or DMEM).

*Axonal length assessment*, αII-tubulin was evaluated. The channel was binarized with the “threshold” tool and then the tool “skeletonize” to convert the signal to a 1 pixel line. With the tool “analyze particles” we could retrieve the total length of the skeletonized signal. After that, total length values were normalized with the values of the vehicle (DMSO or DMEM).

### Differential interference contrast imaging

Differential interference contrast (DIC) images were acquired using a Confocal Olympus FV1200 inverted microscope equipped with a 60x/1.42 NA oil immersion objective and DIC optics (Nomarski prism and polarizer). Transmitted light images were acquired in DIC mode using the Olympus FV10-ASW software. The system uses internal photomultiplier detectors; no external camera was employed. Illumination settings were kept constant across conditions. Image adjustments (brightness and contrast) were applied uniformly in FIJI software (Schindelin et al., 2012) for visualization purposes; no further processing was applied.

### STED nanoscopy

Stimulated Emission Depletion nanoscopy (STED) was performed on two nanoscopes: 1) STED Quad Scan super-resolution microscope (Abberior Instruments, Germany), installed on an Olympus IX83 inverted microscope (Neuro Microscopy Core Facility, McGill University); 2) STED STEDYCON super-resolution microscope (Abberior Instruments, Germany), installed on an Olympus inverted microscope IX81 using a UPlanXApo 100X, NA:1.45, oil immersion objective (CEMINCO, Universidad Nacional de Córdoba). For single color STED, samples were excited with a 635 nm laser, and depleted with a 775 nm laser. For two-color STED, samples were excited with a 580 nm and a 635 nm laser, and depleted with a 775 nm laser. Images were obtained at a single plane, and with the pinhole open to ~1 Airy unit (more details on STED acquisitions using this instrument can be found in the supplier’s website). Briefly, pixel size was set to 10 or 15 nm, and each line was scanned 3 times and accumulated. Confocal excitation laser power was set on a per case basis to avoid saturated pixels in single line scans. During STED, the excitation laser was kept the same as during confocal imaging. Depletion laser power was set to the maximum allowed by the software, to obtain the maximum possible lateral resolution. Imaging conditions were kept across experimental groups belonging to the same experiment, and whole experiments were registered at once.

### MPS organization assessment

#### Sampling of axonal portions to be examined

STED imaging was performed in ROI of ~3 μm x ~3 μm. Selection of these were performed on the confocal signal, with no clue to the underlying nanometer organization. During organization assessment (Autocorrelation or Gollum) 1 μm segments were chosen, aiming at the most representative segment of that ROI. During image analysis, the experimenter was blinded to the group to which each image belonged.

#### Autocorrelation amplitude analyses

in each axon 2 μm segments were chosen and fluorescence intensity profiles were extracted along the longitudinal axis of the axonal segments. These profiles were obtained from high-resolution STED images and subsequently exported to Excel for further analysis. One-dimensional autocorrelation functions were computed from the intensity data to assess spatial periodicity. The mean amplitudes of the first two autocorrelation peaks were quantified to determine the presence and regularity of the expected ~190 nm periodicity associated with the MPS.

#### MPS organization assessment by “Gollum” (Barabas et al., 2017)

this open-source image analysis tool was designed for the automated quantification of the quality of protein periodic structures. Regions of interest were divided in 1 μm × 1 μm sub-regions and analyzed using the Gollum software, which is accessible at https://github.com/cibion-conicet/Gollum. Briefly, the analysis consists of interrogating systematically the presence of a given structure in images, by comparing subregions of the images to a reference pattern. For studying the MPS of MN, we first determined that control axons had a period of ~185 nm and hence the reference pattern consists of gaussian signals with a period of 185 nm. The correlation value obtained corresponds to a two-dimensional Pearson correlation coefficient between the subregion and a modeled MPS period and is indicative of how preserved the periodic organization of the MPS is in that 1 μm section (referred to as *MPS correlation coefficient*). The measurement was repeated along axons in the image with no overlapping, by an operator blind to the experimental conditions.

### Western Blot analyses

For western blot analyses, cells were seeded in 6-well plates. Two-week-old MNs in MN Induction and Maturation Medium were treated with DMSO or 0.1 μM staurosporine for 1 h, then either pelleted immediately or 6 hrs later following a medium change. For protein extraction, cell pellets were lysed in 200 μL of ice-cold 1X RIPA buffer (Millipore, USA) containing a cocktail of protease and phosphatase inhibitors (Roche, Switzerland) and incubated for 30 min at 4℃. Every 10 min, the samples were mechanically disrupted using a pipette. Finally, the samples were centrifuged at 10,000 g for 20 min at 4℃, and supernatants were collected for protein quantification.

Protein concentration in supernatants was quantified using the DC Protein Assay (Bio-Rad, USA) according to the manufacturer’s instructions. For blotting αII-spectrin, βII-spectrin, and SNTF, 10 μg of protein were loaded onto a 7.5% SDS-PAGE gel and run at 70 V for 15 min, then at 120 V for ~1.5 h. Semi-dry transfer to nitrocellulose membranes was performed using the Trans-Blot Turbo Transfer System (Bio-Rad, USA) for 10 min at 2.5 A, up to 25 V. For blotting Caspase-3 and cleaved-Caspase-3, 20 μg of protein were loaded onto a 12% SDS-PAGE gel and run at 70 V for 15 min, then at 120 V for ~1 h. Semi-dry transfer to nitrocellulose membranes was performed using the Trans-Blot Turbo Transfer System for 30 min up to 1.0 A, at 25 V.

After the transfer, membranes were blocked with 5% BSA (or 5% skimmed milk [Bioshop, Canada] for Caspase-3 and cleaved-Caspase-3) in 1X TBS buffer containing 0.1% Tween 20 (blocking solution) for 1 h at RT with continuous shaking, followed by an overnight incubation with primary antibodies, diluted in their respective blocking solutions, at 4℃ with continuous shaking. After three 10 min washes with 1X TBS containing 0.1% Tween 20 (washing solution) and continuous shaking, membranes were incubated with horseradish peroxidase (HRP)-conjugated antibodies, diluted in their respective blocking solutions, for 2 hrs at RT with continuous shaking. After three 10 min washes with washing solution and continuous shaking, protein bands were detected using the ClarityTM Western ECL (Bio-Rad, USA) (or ClarityTM Max Western ECL (Bio-Rad, USA) for cleaved-Caspase-3) according to the manufacturer’s instructions and visualized using a Chemidoc MP Imaging System (Bio-Rad, USA). Quantification was performed with the FIJI software (Schindelin et al., 2012), using β-actin as loading control.

The primary antibodies used were (more details as Supplementary information): anti-βII-spectrin (BD Biosciences), anti-α-II-spectrin (BioLegend), anti-Caspase-3 and anti-Cleaved-Caspase-3 (Cell Signaling) and anti-SNTF (Millipore). And secondary antibodies were: Goat anti-mouse IgG(H+L)-HRP and Goat anti-rabbit IgG(H+L)-HRP (Jackson Immunoresearch).

### Statistical analyses and study design

Data are presented as mean ± SEM and most graphs show also the individual values of the replicates from which the mean and SEM were calculated, as indicated in the figure legends. Calculations were made using the statistical software GraphPad Prism. Normality and homoscedasticity were proved for each group in order to apply the following parametric tests. When comparing two groups, Student’s t test (two-tailed) was used for statistical tests. When comparisons were made among three or more groups, one-way ANOVA was used (or two-way ANOVA was used when more than 1 independent variable was examined); followed by Tukey post-hoc multiple comparisons, to find where the differences occurred. Experiments were performed at least 3 times, using cells from different differentiation batches. Significance is indicated with asterisks when p ≤ 0.05 and detailed in each case. The study design was not pre-registered. No randomization was performed to allocate subjects in the study. No sample size calculation was performed. Outliers were identified, and excluded from analysis, using the ROUT method (Q=1%), implemented through GraphPad Prism.

## Ethics approval

The use of iPSCs in this research was approved by the McGill Research Ethics Board (IRB Study Number A03-M19-22A, titled “The Use of Human Induced Pluripotent Stem Cells (iPSCs) for Modelling Neurodegenerative Disorders”).

## Consent for publication

Not applicable.

## Availability for data and materials

All data generated or analyzed during this study are included in this published article.

## Author contributions

N.G.G., M.J.C.M., T.M.D., and N.U. contributed to the conceptualization of the study by formulating the overarching research goals and aims. The methodology, including the development and design of experimental approaches, was established by N.G.G., M.J.C.M., E.A.G., T.M.D. and N.U. A.K.F., S.L., L.G., G.H., G.M. and T.M.D. produced, validated and provided the iPSC lines. The investigation phase, which included performing experiments and data collection, was undertaken by N.G.G., M.J.C.M., W.E.R. and N.U. Image analysis, data curation and statistical methods were performed by N.G.G., M.J.C.M., G.B., W.E.R. and N.U. Resources, including the provision of materials, instrumentation, and analysis tools, were contributed by F.D.S., A.A., M.B., E.A.G., T.M.D. and N.U. Visualization of data and preparation of figures were carried out by N.G.G., M.J.C.M., and N.U. Project administration, including coordination and execution of the research activities, was led by T.M.D. and N.U. Finally, funding acquisition to support the study was secured by T.M.D., N.U.. The original draft of the manuscript was written by N.G.G., M.J.C.M., T.M.D. and N.U, while all authors contributed to reviewing and editing the final version.

## Acknowledgements

N.G.G. and G.B. have PhD fellowships from CONICET (Consejo Nacional de Investigaciones Científicas y Técnicas). The authors greatly acknowledge the technical and imaging assistance of Dra. Cecilia Sampedro, Dr. Carlos Mas, Dr. Pilar Crespo and Dr. Gonzalo Quassollo from Centro de Micro y Nanoscopía de Córdoba – CEMINCO – CONICET – Universidad Nacional de Córdoba, Córdoba, Argentina. We thank Dr. Thomas Stroh and all members of the Neuro Microscopy Core Facility for their valuable training and assistance with STED and confocal microscopy, which greatly supported this work. Authors want to thank Josefina Alejandra Pogonza Bandrowsky and Dr. Laura Ester Montroull for primary cultures of rat hippocampal neurons.

