## Supplementary information for "BetaII-Spectrin Gaps and Patches Emerge from the Patterned Assembly of the Actin/Spectrin Membrane Skeleton in Human Motor Neuron Axons"

#### Title


### Supplementary Materials and Methods

#### ALS knock-in iPSC lines

Previous studies described the application of a streamlined CRISPR/Cas9 workflow to introduce homozygous knock-in mutations into the AIW002-02 iPSC line, generating iPSC lines with ALS-related mutations: $FUS^{H517Q}$ (Deneault et al. 2022), $SOD1^{G93A}$ and $SOD1^{D90A/G93A}$ (Castellanos-Montiel et al. 2023), and $TDP\text{-}43^{A382T}$ (Lépine et al. 2024). Here, we differentiate these validated iPSC disease lines into MNs to assess changes in the MPS following staurosporine treatment.

#### Animal use and care

Pregnant Wistar rats were born in the vivarium of INIMEC-CONICET-UNC (Córdoba, Argentina). Wistar rats lines were originally provided by Charles River Laboratories International Inc (Wilmington, USA). All procedures and experiments involving animals were approved by the Animal Care and Ethics Committee (CICUAL http://www.institutoferreyra.org/en/cicual-2/) of INIMEC-CONICET-UNC (Resolution numbers 014/2017 B, 015/2017 B, 006/2017 A and 012/2017 A) and were in compliance with approved protocols of the National Institute of Health Guide for the Care and Use of Laboratory Animals (SENASA, Argentina). All methods were carried out in accordance with relevant guidelines and regulations.

#### Primary culture of rat hippocampal neurons and staining

Embryonic day-18 rat embryos (euthanized by CO2 overdose) were used to prepare primary hippocampal cultures as previously described (Bisbal et al., 2018; Wilson et al., 2020a). Briefly, hippocampi from 18 days fetal rats were dissected and incubated with trypsin (0.25% for 15 min at 37°C) (Thermo Fisher Gibco; Cat Number: 15090-046) and mechanically dissociated by trituration with a Pasteur pipette. Cells were plated on Cover Glasses, Circles, 12 mm (Marienfeld Superior; Cat Number: 633029) coated with 1 mg/mL poly-L-lysine (Sigma Chemical Co; Cat Number: P2636) at a density of 2000 cells/cm2 in minimum essential medium (MEM, Thermo Fisher Gibco; Cat Number: 61100-061) supplemented with CTS GlutaMAX I Supplement (Thermo Fisher Gibco; Cat Number: A1286001), Sodium Piruvate (Thermo Fisher Gibco; Cat Number: 11360070), Penicillin-Streptomycin (Thermo Fisher Gibco; Cat Number: 15140122), and 10% horse serum (Thermo Fisher Gibco; Cat Number:16050122). After 2 h, the coverslips were transferred to dishes containing serum-free Neurobasal (Thermo Fisher Gibco; Cat Number: 21103049) with B-27 Plus Supplement (Thermo Fisher Gibco; Cat Number: A3582801) and CTS GlutaMAX I Supplement (Thermo Fisher Gibco; Cat Number: A1286001). Neurons were fixed at 7 DIV in PFS (paraformaldehyde 4%, sucrose 4% in PBS) and stained in standard conditions with antibodies against βII-spectrin, protein 4.1N and βIII-tubulin.

**Table 1. List of reagents for neuronal treatment.**

| Reagent | Working concentration | Solvent (vehicle) | Time of treatment | Reference |
|---|---|---|---|---|

| | | | | |
|---|---|---|---|---|
| Staurosporine | 0.1 μM | DMSO | 1 or 2 hrs | Tocris, cat. #1285 |
| L-glutamate | 0.5 mM | DMEM | 1 hr | Sigma-Aldrich, cat. #G1626 |
| Sodium arsenite | 0.25 μM | DMEM | 1 hr | Sigma-Aldrich, cat. #S7400 |
| Latrunculin A | 0.2, 1, 5 μM | Ethanol | 1 hr or 2 hrs | Cayman, cat. #10010630 |
| ALLN-1 | 5 μM | DMSO | 2 hrs | Cayman, cat. #14921 |
| Caspase-3 inhibitor (zDEVD-fmk) | 50 μM | DMSO | 2 hrs | R&D Systems, cat. #FMK004 |

**Table 2. List of primary and secondary antibodies for immunofluorescence staining**

| **Primary antibodies** | | | |
|---|---|---|---|
| Antibody | Host species | Working dilution | Reference |
| βII-spectrin | Mouse | 1:400 (ICC)<br>1:1000 (WB) | BD Biosciences (cat. #612563) |
| α-II-spectrin | Mouse | 1:400 (ICC)<br>1:1000 (WB) | BioLegend (cat. #803201) |
| βIII-tubulin | Rabbit | 1:1000 (ICC) | BioLegend (cat. #PRB-435P) |
| βIII-tubulin | Chicken | 1:1000 (ICC) | Abcam (cat. #1107216) |
| αII-tubulin | Rat | 1:1000 (ICC) | Sigma-Aldrich (clone α-3A1, cat. #T5168) |
| Caspase-3 | Rabbit | 1:1000 (WB) | Cell Signaling (cat. #9662) |
| Cleaved-Caspase-3 | Rabbit | 1:500 (ICC/WB) | Cell Signaling (cat. #9661) |
| Protein 4.1N | Rabbit | 1:400 (ICC) | Abcam (cat. #ab244499) |
| SNTF | Rabbit | 1:500 (ICC)<br>1:5000 (WB) | Millipore (cat. #ABN2264) |

| | | | |
|---|---|---|---|
| NF-H | Chicken | 1:1000 (ICC) | Abcam (cat. #ab4680) |
| NF-M | Rabbit | 1:1000 (ICC) | Millipore (cat.#AB1987) |
| **Secondary antibodies (ICC)** | | | |
| Goat anti-mouse IgG-STAR ORANGE | | 1:250 | Abberior (cat. #STORAGE-1001) |
| Goat anti-rabbit IgG-STAR ORANGE | | 1:250 | Abberior (cat. #STORAGE-1002) |
| Goat anti-rabbit IgG-STAR RED | | 1:250 | Abberior (cat.#STRED-1002) |
| Goat anti-mouse IgG-Atto 647N | | 1:250 | Millipore (cat. #50185) |
| Donkey anti-chicken IgY-Alexa488 | | 1:500 | Jackson Immunoresearch (cat.#703-545-155) |
| Goat anti-rat IgG-Dylight488 | | 1:500 | Abcam (cat. #ab96887) |
| Phalloidin-Atto 647N | | 1:200 | Sigma-Aldrich (cat. #65906) |

**Table 3. List of secondary antibodies for Western Blot**

| | | |
|---|---|---|
| HRP-conjugated antibodies (WB) | | |
| Goat anti-mouse IgG (H+L) | 1:10000 | Jackson Immunoresearch (cat. #115-035-003) |
| Goat anti-rabbit IgG (H+L) | 1:10000 | Jackson Immunoresearch (cat.111-035-144) |

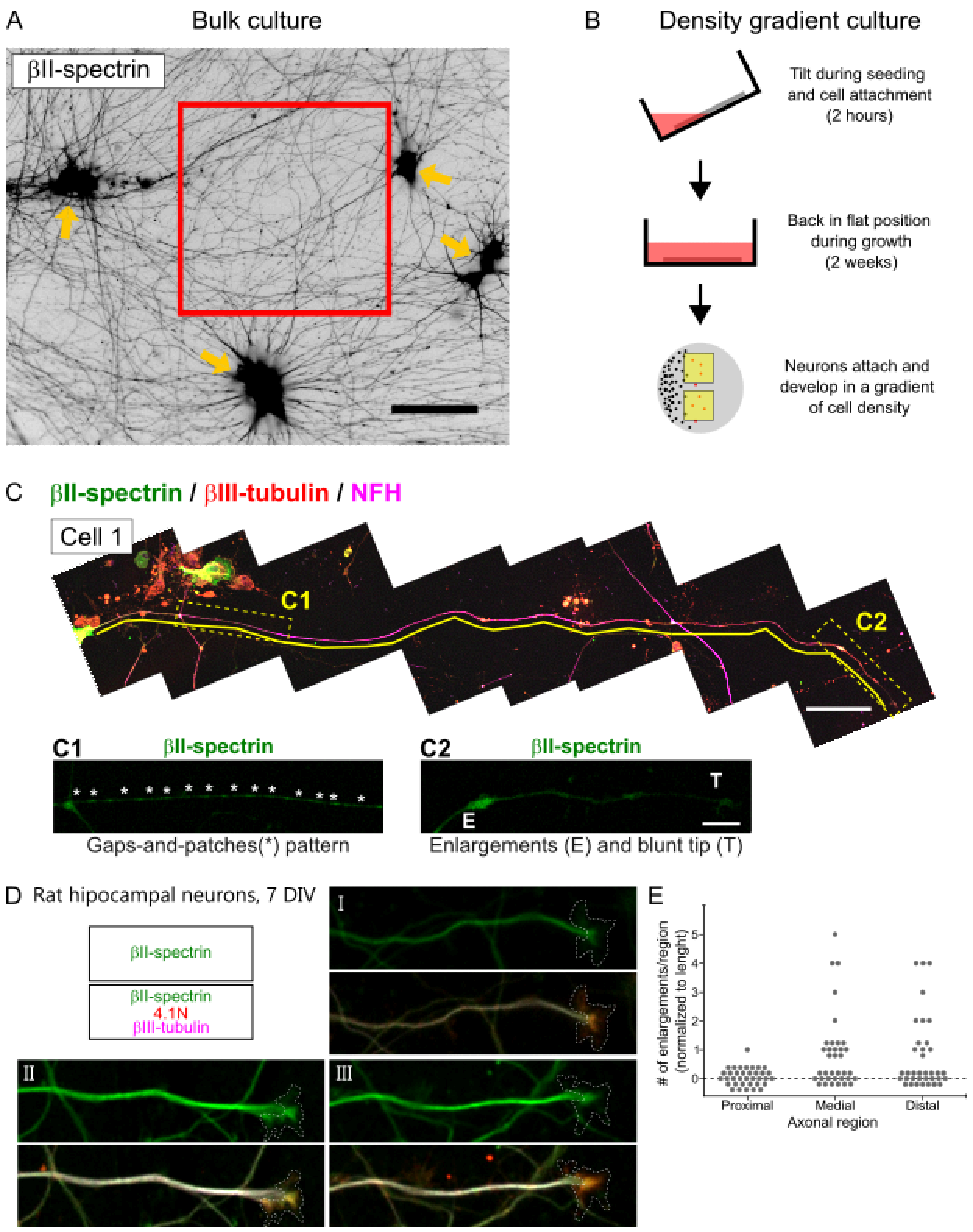


**Supplementary Figure 1.** ***Quantitative and qualitative analyses of βII-spectrin distribution in human iPSC-derived MN axons***

(**A**) Representative confocal image of a 2 weeks old "bulk culture", immunostained for βII-spectrin. Yellow arrows point to groups of >6 neurons. Red square shows the typical sampling performed to assess axonal features. Scale bar = 50 µm. (**B**) Schematic representation of the procedure to generate "density gradient cultures". During seeding and the 2-hour cell attachment period, culture vials containing coverslips were tilted as shown (top). Culture vials were then placed in a standard upright position for cell maintenance (2 weeks for most experiments, middle). For analyses of whole individual axons, areas toward the middle of the coverslip provided a density that supported healthy neurons while allowing the tracing of single axons (yellow squares, bottom). (**C**) Representative reconstruction of an individual axon, immunostained for βII-spectrin, βIII-tubulin and NFH. Yellow line highlights the axon being followed. Scale bar = 50 μm. The C1 dashed rectangle highlights a gap-and-patch pattern (*: patches), whereas the C2 dashed rectangle highlights axonal enlargements (*E*) and the axonal tip (*T*). Scale bar = 10 μm. (**D**) Representative images (I-III) of hippocampal neurons grown for 7 DIV and stained for βII-spectrin, 4.1N and βIII-tubulin. The dashed line contours the expanded growth cone at the tip of the axon, which was delineated using the actin-binding protein 4.1N. Scale bar = 10 μm. (**E**) Quantification of number of axonal enlargements in proximal, medial and distal sections.

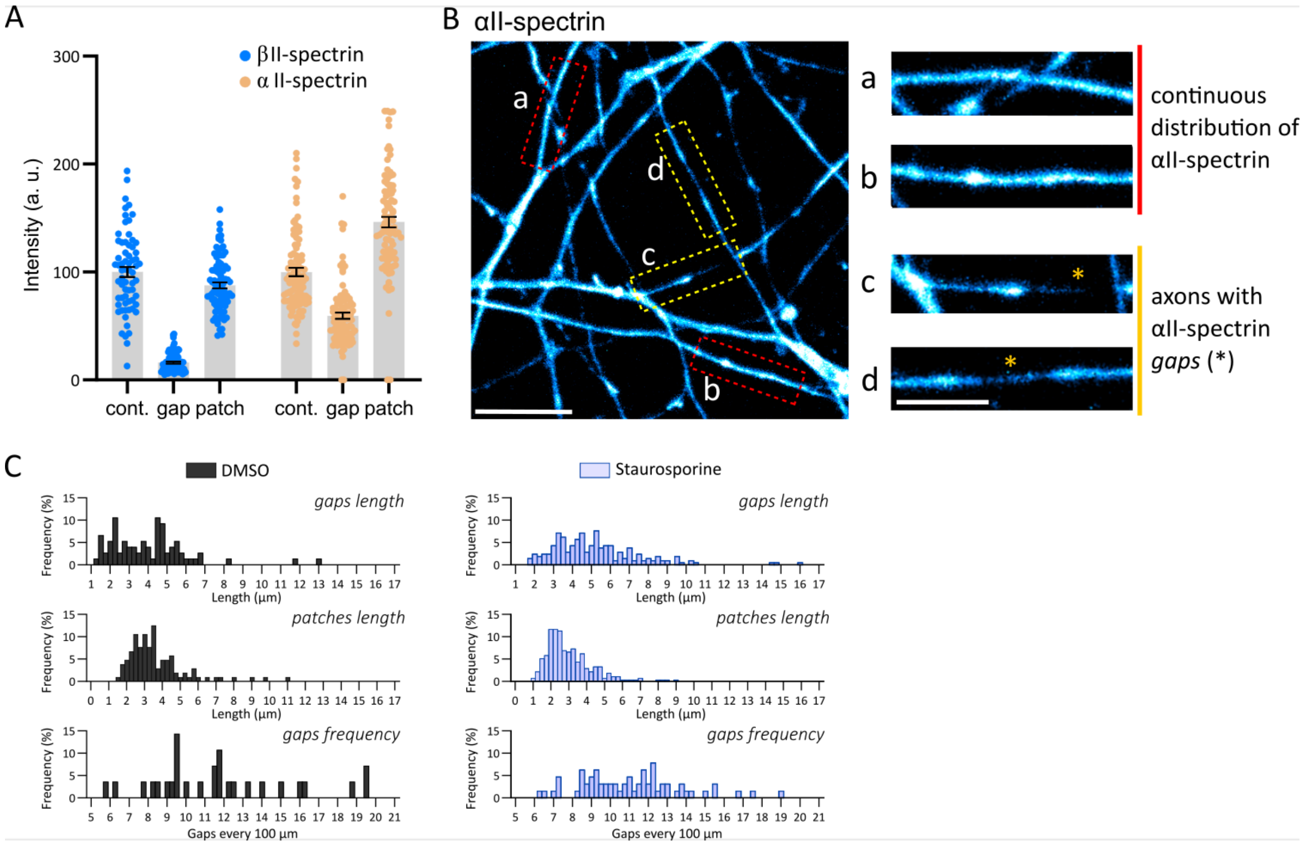


**Supplementary Figure 2. *Axons of human iPSC-derived motor neurons present interruptions in αII-spectrin.***

(**A**) Intensity (mean grey values, a.u.) of βII-spectrin and αII-spectrin within regions with continuous distribution (cont.), within gaps and within patches. (**B**) Confocal images of bulk cultures immunostained for αII-spectrin. Inserts a-c show examples of axons with a continuous distribution of βII-spectrin (a and b) and axons with sharp interruptions or "gaps" (asterisks in c and d). Scale bar = 10 µm (left panel) and 5 µm (zoom-in inserts). (**C**) Histograms showing percentual frequency of gaps length, patches length and gap frequency per 100 µm, in control (DMSO) and staurosporine treated cells.

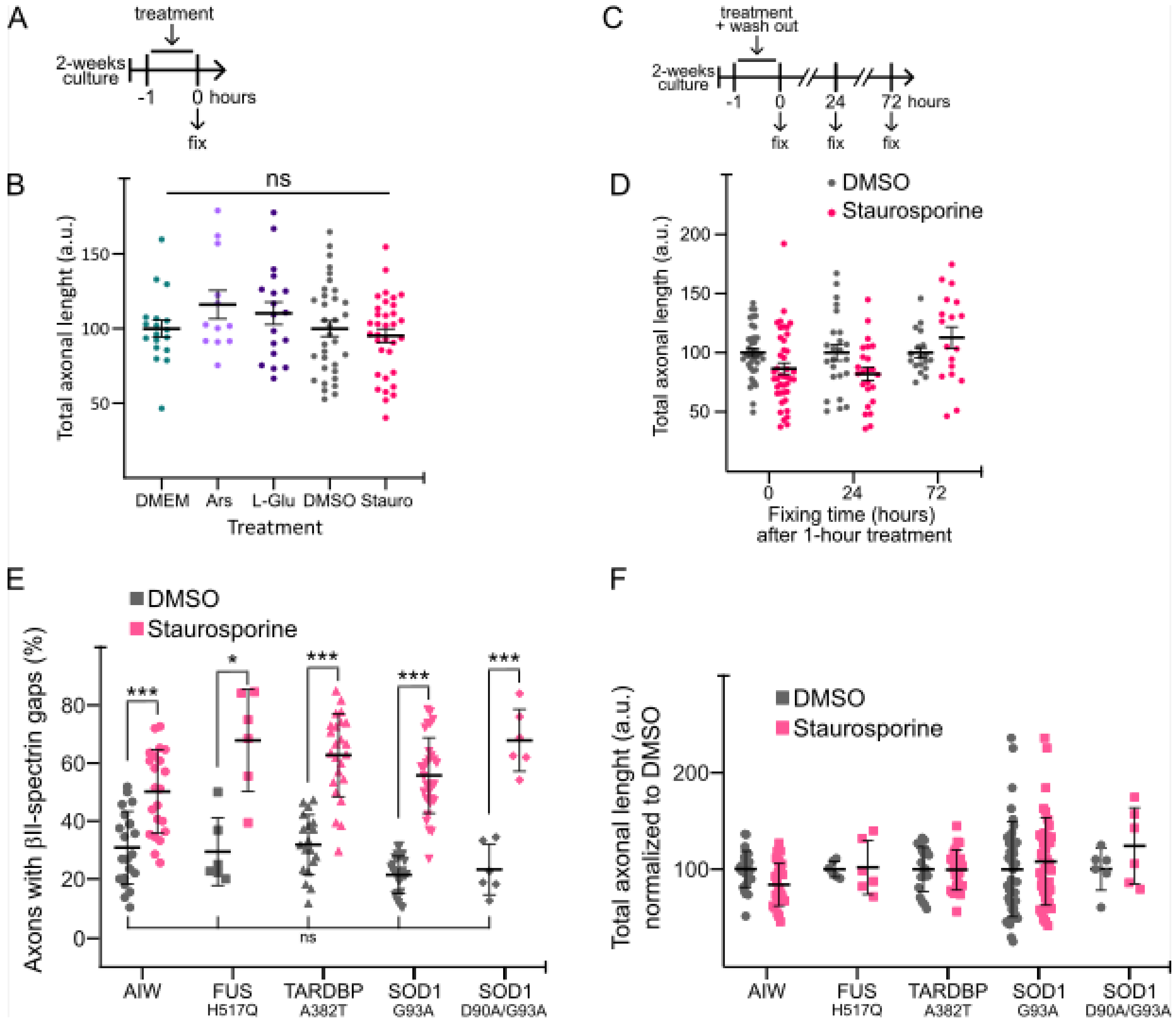


**Supplementary Figure 3. Interruptions in βII-spectrin increase as a function of weeks in culture and by acute treatment with staurosporine.**

(**A**) Schematic representation of the timeline of the experiments shown in Fig. 3B. (**B**) Total axonal length (a.u.; based on α-tubulin staining) of 2-week-old MNs treated for 1 h with low doses of arsenite (2 μM), L-glutamate (0.05 mM) and staurosporine (0.1 μM), or the control vehicles. ns: not significant. (**C**) Schematic representation of the timeline of the experiments shown in Fig. 3C. (**D**) Total axonal length (a.u.; based on α-tubulin staining) of 2-week-old MNs treated for 1 h with DMSO or staurosporine (0.1μM) and fixed immediately (0), or 24 and 72 hrs later. (**E-F**). Percentage of axons with βII-spectrin gaps (E) and total axonal length (F, a.u.; normalized to DMSO treatment in each line) of 2-week-old MNs treated for 1 h with DMSO or staurosporine (0.1 μM) and fixed immediately, across the control line (AIW) and lines carrying mutations in ALS-related genes: FUS (H517Q), TDP43/TARDBP (A382T) and SOD1 (G93A and D90A/G93A). *: $p<0.05$; ***: $p<0.001$.

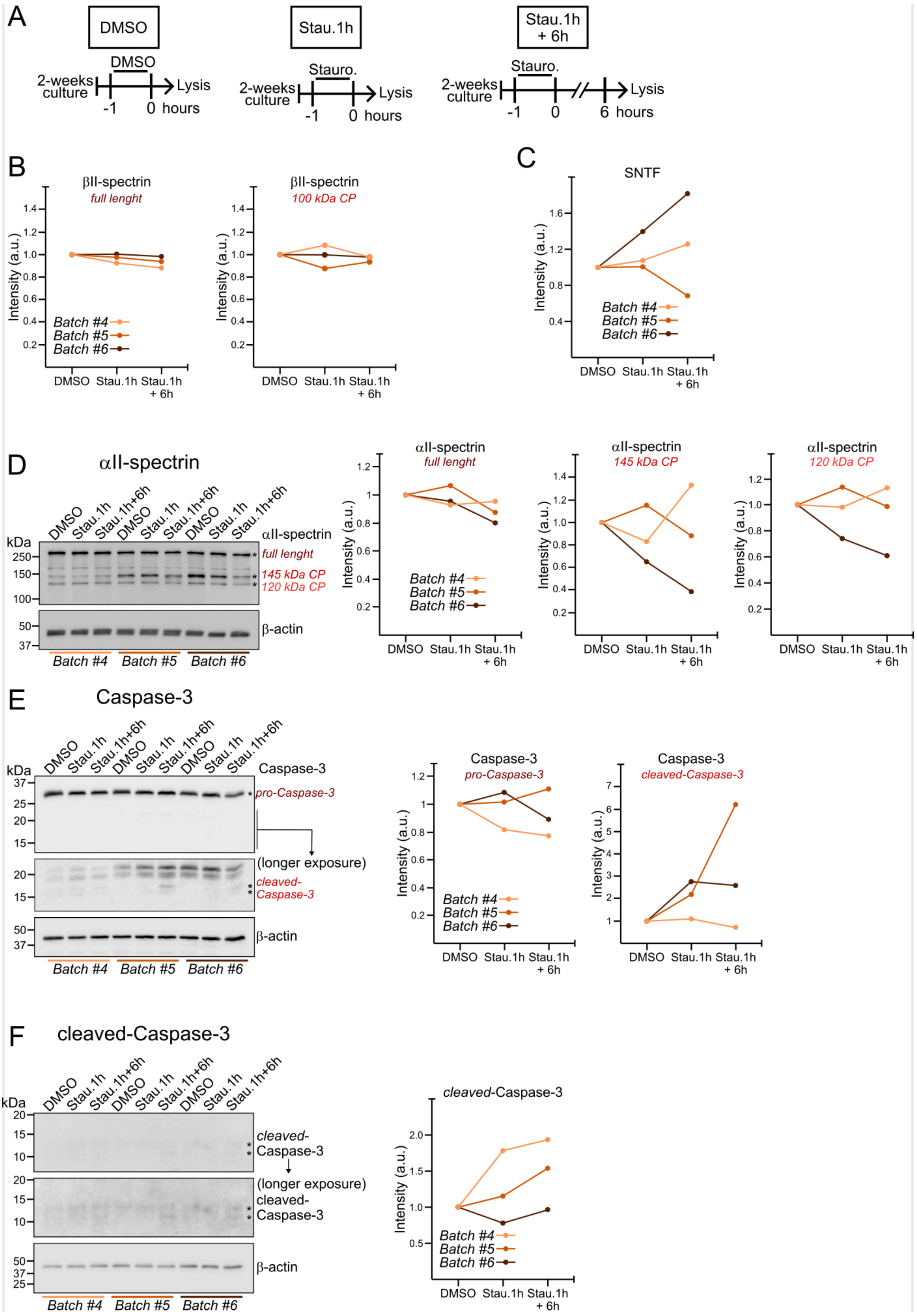
A
DMSO
Stau.1h
Stau.1h + 6h
2-weeks culture
Stauro.
Lysis
-1
0
6
hours
B
βII-spectrin
full lenght
βII-spectrin
100 kDa CP
Intensity (a.u.)
Batch #4
Batch #5
Batch #6
DMSO
Stau.1h
Stau.1h + 6h
C
SNTF
D
αII-spectrin
145 kDa CP
120 kDa CP
kDa
β-actin
E
Caspase-3
pro-Caspase-3
cleaved-Caspase-3
(longer exposure)
F
cleaved-Caspase-3

**Supplementary Figure 4.** ***Proteolytic cleavage of spectrins is not a mechanism that forms gaps in the MPS lattice.***

(**A**) Schematic representation of the experimental design and treatment timeline analyzed by western blot (results shown in Figure 4 and in this figure). (**B**) Quantification of the densitometry of βII-spectrin reactive bands indicated in Fig. 4A (*). (**C**) Quantification of the densitometry of SNTF (αII-Spectrin N-Terminal Fragment) reactive bands indicated in Fig. 4B (*). (**D**) Western blot image against αII-spectrin and β-actin (left) and quantification of the densitometry of reactive bands (*, right). (**E**) Western blot image against Caspase-3 and β-actin (left) and quantification of the densitometry of reactive bands (*, right). (**F**) Western blot image against cleaved-Caspase-3 and β-actin (left) and quantification of the densitometry of reactive bands (*, right).

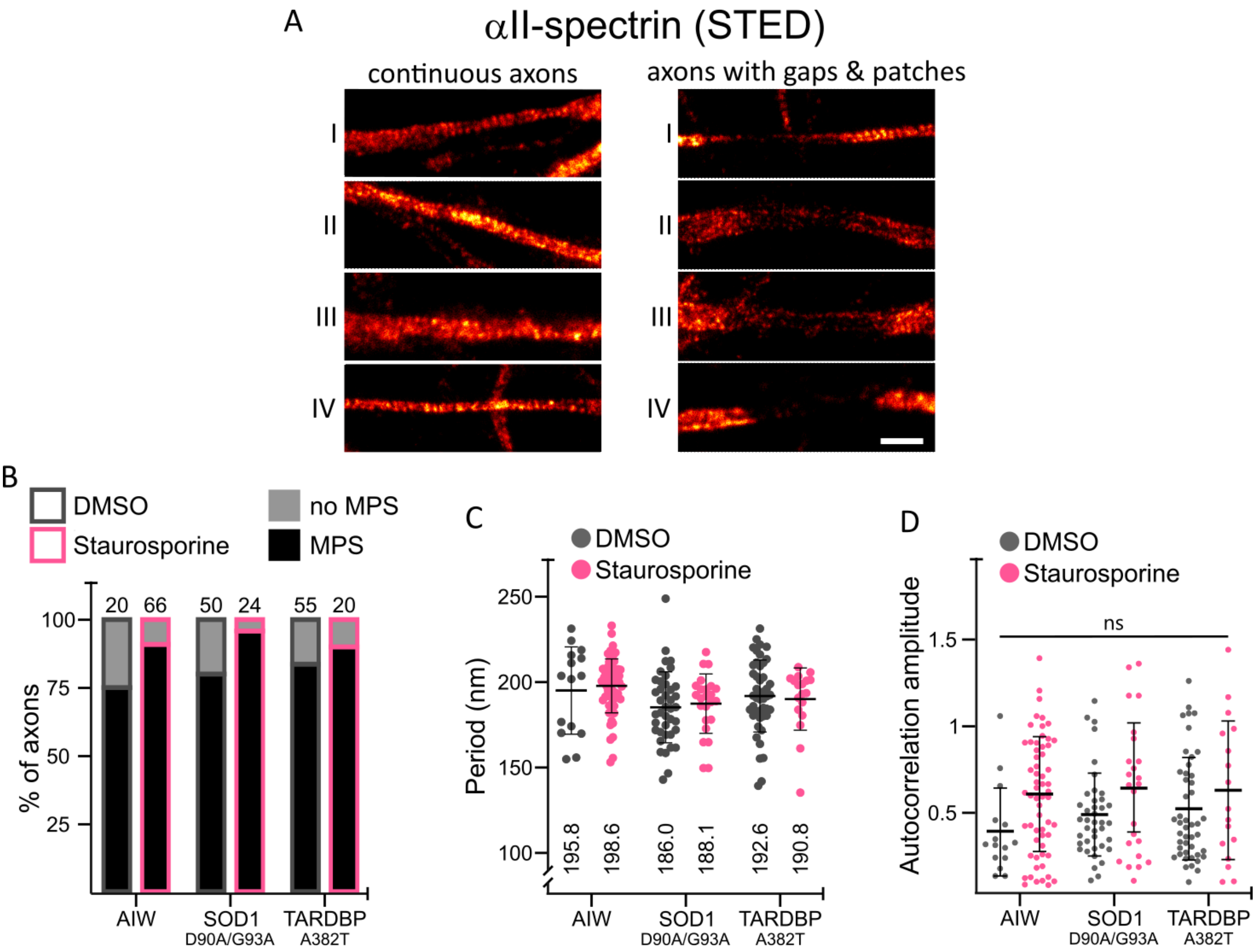


**Supplementary Figure 5. The MPS is absent in βII-spectrin gaps, but it is well organized within patches**

(**A**) Representative STED images (of αII-spectrin acquired using STED microscopy in axons with continuous αII-spectrin distribution (I-IV) and in axons with αII-spectrin-gaps (V-VIII). Scale bar = 1 µm.

(B) Analyses of the percentage of axonal sections that presented a MPS in 2-week-old MNs treated for 1 h with DMSO or staurosporine (0.1 µM) and fixed immediately, across the control line (AIW) and lines carrying mutations in ALS-related genes: SOD1 (D90A/G93A) and TDP43/TARDBP (A382T). Above each bar, the total number of sections analysed per group is shown.

(C-D) Period (C) and autocorrelation amplitude values (D) of MPS axonal sections shown in B. In (C) values inserted in the graph are the mean value of each group.

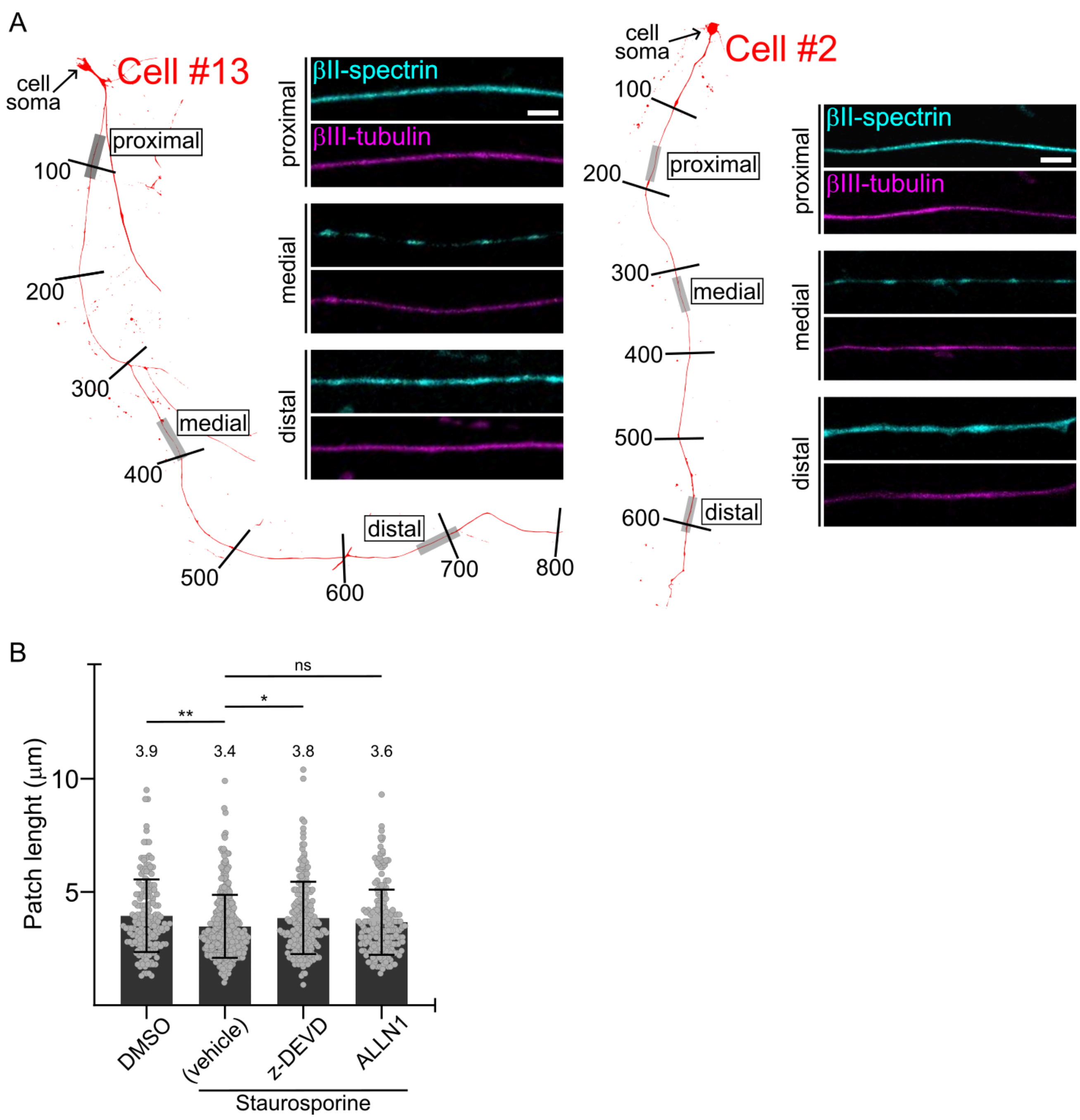


**Supplementary Figure 6. Gaps and patches preferentially occur in the middle section of axons**
(**A**) Representative axonal reconstructions of two neurons. Numbers along axons indicate approximate distance to the cell soma (µm). Inserts show confocal images of proximal, medial and distal regions, indicated as grey rectangles in the axonal reconstruction, immunostained for βII-spectrin (cyan) and βIII-tubulin (magenta). Scale bar = 1 µm.
(B) Quantification of patches length found in vehicle-treated axons (DMSO) and staurosporine-treated 2-week-old MNs with or without the inhibitors for caspase-3 (z-DEVD-fmk) or calpains (ALLN1). In the graph, the values above each bar is the mean length (µm) of each group. One way ANOVA, *: $p<0.05$, **: $p<0.01$.
